# The oncogene SLC35F2 is a high-specificity transporter for the micronutrients queuine and queuosine

**DOI:** 10.1101/2024.12.03.625470

**Authors:** Lyubomyr Burtnyak, Yifeng Yuan, Xiaobei Pan, Lankani Gunaratne, Gabriel Silveira d’Almeida, Maria Martinelli, Colbie Reed, Jessie Fernandez Garcia, Bhargesh Indravadan Patel, Isaac Marquez, Ann E. Ehrenhofer-Murray, Manal A. Swairjo, Juan D. Alfonzo, Brian D. Green, Vincent P. Kelly, Valérie de Crécy-Lagard

**Author notes:** **Corresponding authors:** Valérie de Crécy-Lagard and Vincent P. Kelly **Email:**. eSTEAMed Learning Inc., Maitland, FL 32751, USA. Data Science Institute, Medical College of Wisconsin, Milwaukee, Wisconsin 53226, USA. **Author Contributions:** Conceptualization, VdC-L, VPK; methodology, VdL, VPK, JA, MAS, AEM; formal analysis VdC-L, VPK; investigation, LB, YY, XP, BDG, LG,GSdA, MM, JFG, CR. BIP, IM; writing original draft, YY, LB, AEM, VdC-L and VPK.; funding acquisition, VdC-L, VPK, MAS, JA, BDG, AEM. **Competing Interest Statement:** The authors declare no competing interests.

## Abstract

The nucleobase queuine (q) and its nucleoside queuosine (Q) are micronutrients derived from bacteria that are acquired from the gut microbiome and/or diet in humans. Following cellular uptake, Q is incorporated at the wobble base (position 34) of tRNAs with a GUN anticodon, which is important for efficient translation. Early studies suggested that cytosolic uptake of queuine is mediated by a selective transporter that is regulated by mitogenic signals, but the identity of this transporter has remained elusive. Here, through a cross-species bioinformatic search and genetic validation, we have identified the solute carrier family member SLC35F2 as a unique transporter for both queuine and queuosine in *Schizosaccharomyces pombe* and *Trypanosoma brucei*. Furthermore, gene disruption in HeLa cells revealed that SLC35F2 is the sole transporter for queuosine in HeLa cells (K_m_ 174 nM) and a high-affinity transporter for the queuine nucleobase (K_m_ 67 nM), with the presence of another low-affinity transporter (K_m_ 259 nM) in these cells. Competition uptake studies show that SLC35F2 is not a general transporter for other canonical ribonucleobases or ribonucleosides, but selectively imports q and Q. The identification of SLC35F2, an oncogene, as the transporter of both q and Q advances our understanding of how intracellular levels of queuine and queuosine are regulated and how their deficiency contributes to a variety of pathophysiological conditions, including neurological disorders and cancer.

**Significance Statement:** The discovery of SLC35F2 as the eukaryotic transporter of queuine and queuosine is key to understanding how these micronutrients are salvaged from the human gut and distributed to different body tissues. Queuosine modification of tRNAs enhances the accuracy and efficiency of codon-anticodon pairing and regulates a range of biological and pathophysiological states, including oxidative stress responses, cancer, learning, memory, and gut homeostasis.

## Introduction

Human health is deeply intertwined with the acquisition of sufficient daily nutrients. This encompasses not only adequate caloric intake, but also a diverse array of essential vitamins, minerals, and other micronutrients necessary for optimal cellular function, disease prevention, and overall well-being (1). Queuine (q) and its nucleoside derivative queuosine (Q) are one such group of micronutrients that are synthesized exclusively by bacteria and salvaged by eukaryotic organisms including plants, fungi, yeast, and metazoan species, with q undergoing subsequent incorporation into tRNAs decoding His, Tyr, Asp, and Asn codons and serving as a substrate for further glycosylation, turnover, and salvage (2) (Fig. 1). Currently, no transporters for intracellular uptake of Q nor q have been identified in any eukaryote and, in metazoans, the mechanism of uptake from the gut and redistribution throughout the body is poorly understood.

**Figure 1.**
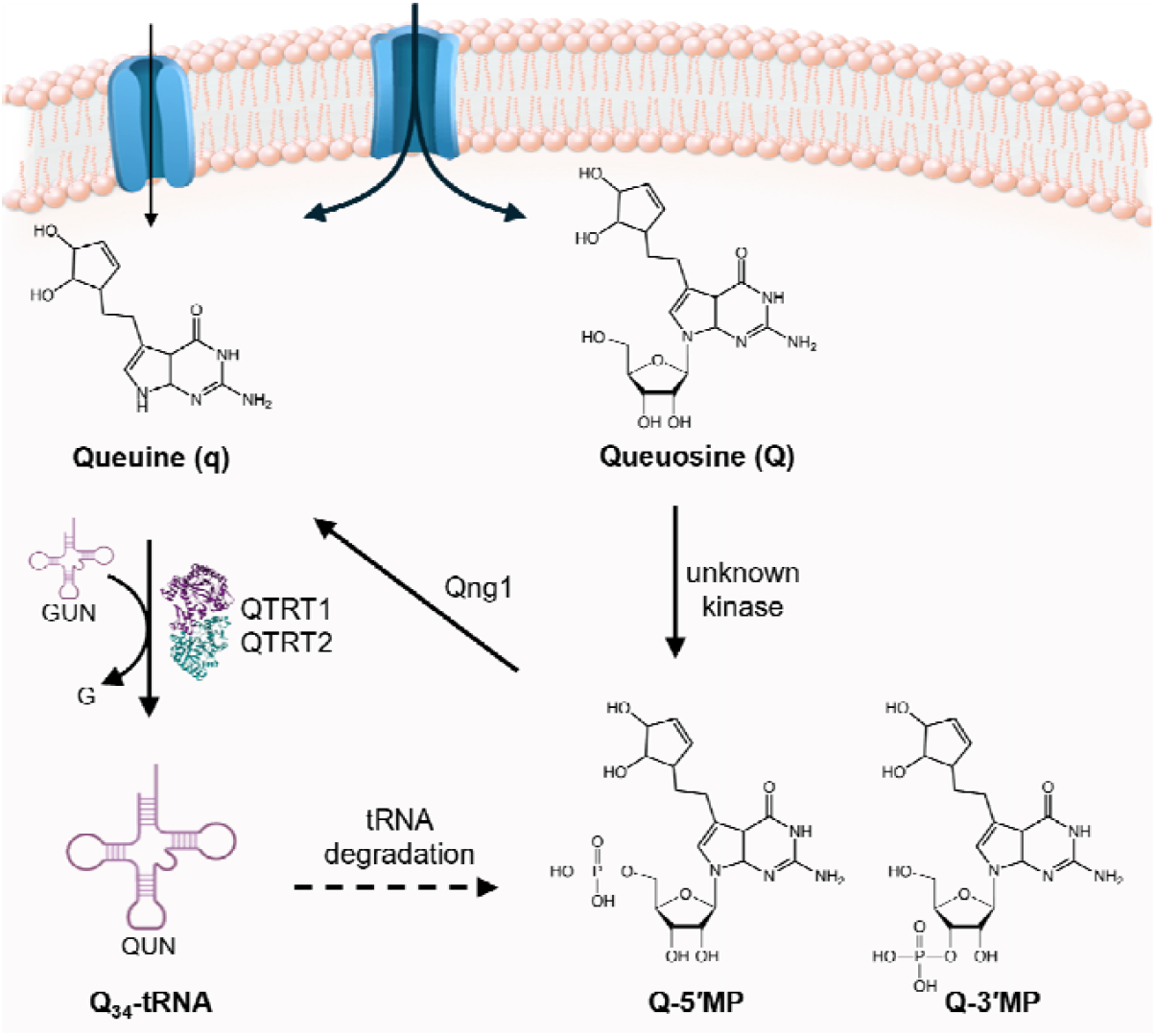
Transport of q and Q into eukaryotes and subsequent intracellular utilization. The q/Q micronutrients are imported into the cell via (an) unidentified transporter(s). Q is phosphorylated by an unknown kinase to Q-5′-monophosphate (Q-5′MP) and subsequently hydrolyzed by QNG1 to release the q nucleobase. Queuine serves as a substrate of the eukaryotic tRNA-guanine transglycosylase (TGT) heterodimer (QTRT1 and QTRT2) for incorporation into tRNA of the GUN anticodon, generating tRNAs modified with Q (Q_34_-tRNA) at position 34. The Q in tyrosyl- and aspartyl-tRNA can be modified further by galactosylation and mannosylation by the enzymes QTMAN and QTGAL, respectively (61). Q-5′MP is the biologically relevant substrate of QNG1, a queuosine-5′-monophosphate hydrolase (18).

In humans, the circulating levels of q are estimated to be in the range of 1-10 nM (3) recent studies indicating a mean serum concentration of 8 nM in females and 6.8 nM in males (4). Early studies in fibroblasts and mammalian cell lines showed that a specific cellular uptake mechanism exists to transport q from the extracellular space to the cytosol. Using a tritiated q derivative (rQT_3_), it was shown that uptake in human foreskin fibroblasts can occur *via* two different transporters, a low K_m_ transporter (30 nM) that reached saturation after 2 minutes (13.5 pmol/10^6^ cells/h) and a slower, high K_m_ mechanism (350 nM) that displayed a linear uptake after 3-4 hrs (2.3 pmol/10^6^ cells/h). Moreover, competitive uptake experiments with a variety of purine and purine-derivatives did not affect q transport, suggesting that the transporter is specific to q (5).

Earlier studies indicated that mitogenic signalling pathways could modulate q transport (6). In fibroblasts, low doses and short-term exposure to the Protein Kinase C (PKC) activator phorbol-12, 13-didecanoate (PDD) led to increased rQT_3_ uptake, whereas extended exposure durations inhibited uptake. In addition, the protein phosphatase inhibitors calyculin A and okadaic acid were found to increase rQT_3_ uptake but avoidance of excess exposure time and concentrations were necessary [6]. Later, other PKC activators, including phosphatidylserine, calcium ionophore and diolein were also found to increase rQT_3_ uptake (7). Furthermore, growth factors such as epidermal growth factor, fibroblast growth factor and platelet-derived growth factor were also shown to increase q uptake, both alone and in combination with PKC activators (7). Notably, although a major portion of q transport appears to be regulated by PKC and mitogenic signalling, a substantial basal rate of q uptake exists, which operates independently of exogenous signalling. This implies the presence of two or more transporters for q uptake, one of which may be PKC- and mitogen-regulated.

Here we show that SLC35F2 functions as the exclusive transporter for q and Q in *Schizosaccharomyces pombe* and the human parasite *Trypanosoma brucei*. In the human HeLa cell line, it serves as the primary high-affinity transporter for q and as the unique transporter for Q. Competitive uptake studies reveal that SLC35F2 does not recognize canonical nucleobases nor nucleosides, since they were unable to disrupt q/Q import into HeLa cells. Defining the role of the SLC35F2 transporter, an identified oncogene, as the q/Q transporter has significance for understanding the uptake, distribution and function of this micronutrient in eukaryotes.

## Results

### Prediction of a candidate Q/q transporter family by comparative genomics

The Q modification of tRNA has been independently lost in several eukaryotic clades including the model organism *Saccharomyces cerevisiae*, while being present in most sequenced fungal genomes including *S. pombe* (8). The presence/absence of Q can be inferred by the presence/absence of the signature gene *Qtrt1*, which encodes tRNA guanine transglycosylase that exchanges G for Q at position 34 in tRNA. Previous phyletic profile searches to identify the missing q/Q transporter gene(s) by searching for protein families that co-distribute with *Qtrt1* were performed across three eukaryotic kingdoms (fungi, viridiplantae and metazoa) (8). However, these searches only returned candidate genes for the already characterized *Qtrt1* and *Qng1* genes, the latter being responsible for the release of q from the Q precursor queuosine 5′-monophosphate (Q-5′MP). As an alternate strategy, we analyzed protein families containing transmembrane domains separately in four different taxonomic clades (Sar, Viridiplantae, Metazoa and Fungi), each including around 200 organisms encoding QTRT1 (QTRT^+^) and at least 10 organisms that are QTRT1 deficient (QTRT^-^) (Fig. S1 Step I). The proteins in each of these taxonomic groups containing at least four transmembrane (TM) domains covering 25% or more of their length were clustered based on sequence similarity as described in the methods section (Fig. S1 Step II). The choice of four TM domains was because 96.4% of all known electrochemical potential-driven transporters have at least four TM domains (9). Each cluster within a given group was given a probability score that would be 1 if all the proteins of the cluster were encoded by QTRT^+^ organisms and absent in QTRT^-^ organisms as described in the methods section and in Fig. S1 (Step III).

The top candidate in the fungi clade, which also scored highly in the other three analyzed taxonomic groups, was the IPR052221 annotated in InterPro database as the “SLC35F solute transporter family” (Table S1), but corresponds to the “solute carrier family 35 member F1/2” or the “SLC35F1/F2” KEGG orthology group (K15287)(10). This family of transporters contains 10 transmembrane helices (Fig. S1A) and is a subgroup of the SLC35 superfamily (11). Most members of this family transport nucleoside di-phosphate (NDP) sugars, but a few transport vitamins such as thiamin (12). To date, the substrate specificity of the SLC35F1/F2 subgroup is not known (13).

SLC35F1/F2 proteins were present in 184 out of 189 QTRT^+^ fungi and absent in all of 189 QTRT^-^ Fungi. The gene encoding a SLC35F protein in *S. pombe, SPCC320*.*08*, is co-expressed with ribosome biogenesis genes (https://coxpresdb.jp/locus/?gene_id=2539011). Another published co-expression analysis along the *S. pombe* cell cycle (14) found that *SPCC320*.*08* was the only transporter gene in a cluster of co-expressed genes highly enriched (21 out of 24, Table S2) in rRNA processing and translation related proteins.

We constructed a sequence similarity network (SSN) of the IPR052221 family proteins to dissect their functions (Fig. 2A). The Chordata F1 and F2 paralogous subgroups could not be separated with an alignment score threshold (AST) of 110 (Fig. 2A), but the divergence becomes evident with an AST of 125 (Fig. 2B), indicating similar functions. Members of this family separate mostly following taxonomic lineages except for the Viridiplantae SLC35F proteins, which are split in two groups: one that contains members from QTRT^-^ organisms, such as *Arabidopsis thaliana* (**Table S3**), and one that lacks *A. thaliana* homologs but contains members only from QTRT^+^ organisms, such as *Zea mays*. Indeed, the *Z. mays* genome encodes the full Q salvage pathway (Uniprot accession: QTRT1, A0A3L6E559; QTRT2, A0A8J8Y6E7 and Qng1, A0A3L6FZJ1) while *A. thaliana* is known to lack Q in tRNA, as this wobble base modification has been lost in cruciferous plants (8). The identification of distinct Viridiplantae paralogous groups, one with no *A. thaliana* member, suggested that the SLC35F family is not isofunctional and could also explain why this transporter family was not identified in previous phylogenetic distribution searches that were filtering for transporter families present in *Zea mays* and absent in *A. thaliana* (8). Analyses of the co-expression data available for the two human SLC35F genes link SLC35F2 to ribosome biogenesis and RNA processing, but associate SLC35F1 to neural and synaptic function (Fig. S2). In addition, overexpression of SLC35F1 in HeLa cells recently showed that the transporter resides exclusively on recycling endosomes (15). This would suggest that SLC35F1 is likely not the plasma membrane transporter for q/Q. The combination of *in silico-derived* evidence (presence/absence correlated with QTRT1, co-expression with RNA processing proteins and membrane protein topology) led us to test the hypothesis that SLC35F2 is one of the missing human Q/q transporters.

**Figure 2.**
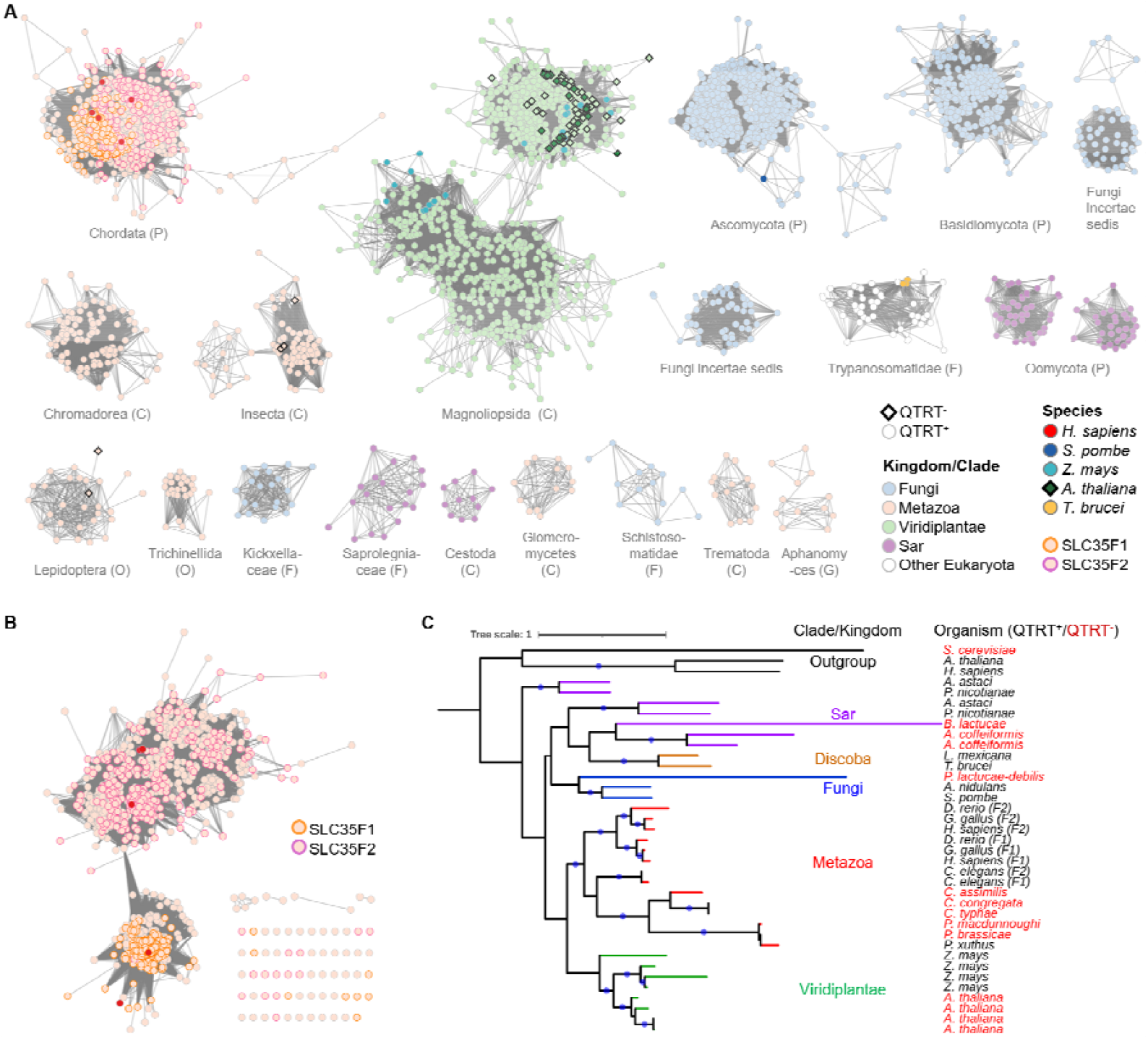
The SLC35F transporter family is functionally diverse. (A) The SSN of SLC35F family proteins in Eukaryota. Each node in the network represents one SLC35F protein. An edge (represented as a line) is drawn between two nodes with a BLAST E-value cutoff of better than 10^−110^ (alignment score threshold of 110). The shape of nodes is based on the presence (circles) and absence (diamonds) of QTRT1 in the organism. The nodes of selected model organisms are highlighted in different colors. The borders of SLC35F1 and F2 annotated in Metazoa are highlighted in orange and pink, respectively. For better visualization, clusters with less than ten sequences are hidden. (B) The sub-SSN of the cluster of phylum Chordata. An edge (represented as a line) is drawn between two nodes with a BLAST E-value cutoff of better than 10-125 (alignment score threshold of 125). (C) The maximum likelihood tree of 34 SLC35F proteins. SLC35B1-4/HUT1 proteins were used as the outgroup. The branches are colored by taxonomic ranks and branches with a 0.7 or more bootstrap value are indicated by blue dots. The QTRT^+^ and QTRT^-^ organisms are shown in black and red, respectively.

### The human *SLC35F2* gene encodes the Q/q transporter

To investigate whether the SLC35F2 protein is required for q or Q transport in humans, a knockout approach was adopted. The SLC35F2 gene is located on the reverse strand of chromosome 11, comprising eight exons that encode a 374-amino acid protein. CRISPR-Cas9 was used to insert a promoter trap into exon two of the gene by homologous recombination — providing puromycin resistance under the control of the SLC35F2 promoter (Fig. S3A). From the clones surviving puromycin treatment, four were selected and genotyped. All four clones were homozygous knockouts for SLC35F2, with either no PCR amplicon observed for the wild-type allele or a band of the incorrect size (Fig. S3B). Strikingly, 3-AcrylamidoPhenylBoronic acid (APB) Northern blot analysis, which can separate Q modified from unmodified tRNAs (16), revealed that in the absence of SLC35F2, all four HeLa cell lines were devoid of detectable Q modification in tRNA^His^ after incubation with Q (25 nM), contrasting with wildtype cells, where approximately fifty percent of the tRNA was Q-modified (Fig. 3A, upper panel). In the case of q nucleobase (25 nM), in wild-type cells, all the tRNA was found to be fully Q-modified with the addition of the micronutrient at a concentration of 25 nM, but a dramatic reduction in modification was observed in the SLC35F2 knockout clones (Fig. 3A, lower panel). To validate and extend this finding, Clone 1 was selected for further analysis by treating cells with increasing concentrations (25 to 250 nM) of either the q-base or Q-nucleoside (Fig. 3B). Notably, at all concentrations tested Q failed to modify the tRNA in the SLC35F2 knockout cells (upper panel), whereas in the case of q, almost a ten-fold higher concentration was required, relative to wild-type cells, to completely modify the tRNA (lower panel). These data show that SLC35F2 is a unique transporter for Q-nucleoside in human cells and the primary high-affinity transporter for the uptake of q base.

**Figure 3.**
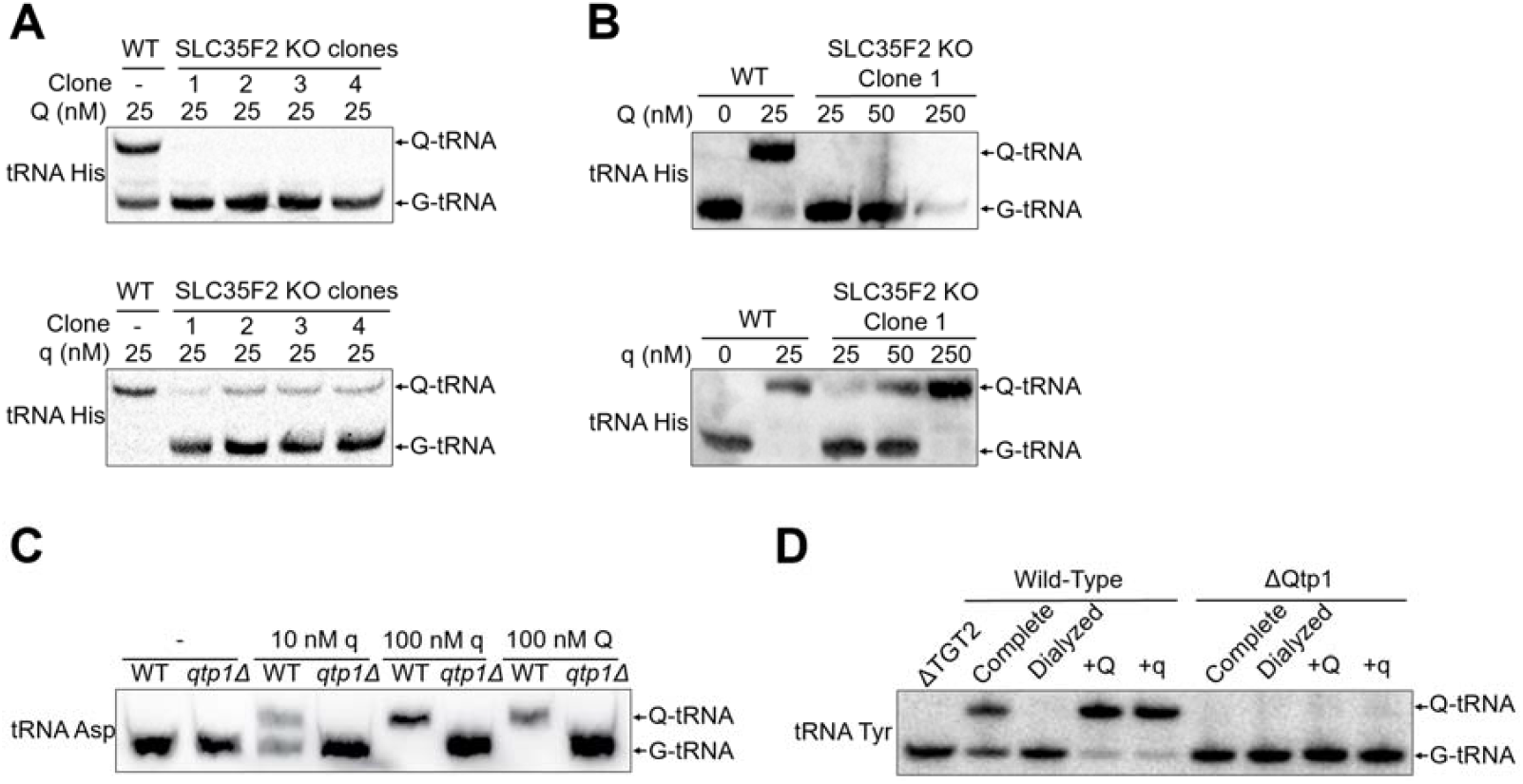
SLC35F2 is the transmembrane transporter responsible for Q/q uptake in humans, *S. pombe* and *T. brucei*. **(A)** tRNA^His^ is Q-modified in wild-type (WT) HeLa cells, but not in four SLC35F2 knockout candidate clones (1-4), as determined by APB Northern blot. Cells were incubated with Q (25 nM) or q (25 nM) for 6 hrs; **(B)** SLC35F2 knock-out clone 1 was incubated with increasing concentrations of q and Q as indicated for 6 hrs, and the Q-modification status of tRNA^His^ was analyzed; **(C)** WT and a *qtp1*Δ *S. pombe* strain were cultured in the absence (-) or presence of q/Q at the indicated concentrations, tRNA extracted and analyzed by APB Northern using a tRNA^Asp^ probe; **(D)** APB Northern analysis of Q-modification status of tRNA^Tyr^ in WT and Δ*qtp1 T. brucei* strains that were cultured in media supplemented with either complete or dialyzed Fetal Bovine Serum (FBS) supplemented when noted with 25 nM Q or q. tRNA extracted from a strain deleted for the gene encoding TGT2, one paralog of TGT in *T. brucei* (21).

### The SLC35F2 homologs of *T. brucei* and *S. pombe* are the sole q/Q transporters in these organisms

Having shown that SLC35F2 was required for both q and Q transport in human cells, we investigated if this was also the case in eukaryotes that encode only one SLC35F protein (Fig. 3C-D). We focused on two single-cell eukaryotes where Q synthesis and function have been extensively studied: the fission yeast *S. pombe* (8, 17–20) and the parasite *T. brucei* (21–23).

The fission yeast *S. pombe* encodes an SLC35F1/F2 homolog SPCC320.08, (here renamed Qtp1 [**Q**ueuosine and queuine **t**rans**p**orter 1]), which has no apparent homolog in *Saccharomyces cerevisiae* (PomBase,(24)). To test whether Qtp1 transports q and/or Q in *S. pombe*, a *qtp1*Δ strain was constructed, and Q levels analyzed via APB Northerns in tRNA^Asp^ extracted from cells grown in medium supplemented with either q or Q (**Fig. 3C**). In the presence of 10 nM q, the wildtype cells showed partial Q modification of tRNA^Asp^, and 100 nM q resulted in full modification, as determined by APB Northern blotting. In contrast, in the *qtp1*Δ strain, tRNA^Asp^ showed no Q modification of the tRNA even at the higher q concentration. Similarly, the *qtp1*Δ strain showed no Q modification of tRNA^Asp^ when provided with the Q nucleoside, whereas the wildtype strain showed full modification.

To study the function of the SLC35F2 homolog in *T. brucei* (ORF Tb927.4.320), here renamed Qtp1 as above for *S. pombe*, both alleles of the corresponding gene were deleted using the CRISPR/Cas9 system (Fig. S4). This gene was not essential in *T. brucei* and growth was not affected (Fig. S5) in agreement with the fact that Q in tRNA is not essential for growth in this organism (21–23). We then extracted total RNA from wild type and knockout cells lines and analyzed Q-content in tRNA^Tyr^ via APB Northern blotting (Fig. 3D). For this analysis, Q-content from cells grown in complete media (non-dialyzed serum) were compared to cells grown in media containing dialyzed serum. No Q-tRNA was detected in cells grown with dialyzed serum, regardless of the presence or absence of Qtp1. This is expected given that serum is the only source of q/Q for *T. brucei* when grown in culture. Similar experiments were then performed by supplementing the dialyzed serum with q or Q (at 25 nM concentration, respectively). In this case, only wild-type cells showed detectable Q-tRNA, while Q-tRNA was undetectable in the *qtp1* knockout (Fig. 3D) even when the cultures were supplemented with 250 nM q (Fig. S6) or when the knockout cells were grown in complete media (**Fig. 3D**).

Taken together, these experiments show that in two organisms encoding only one member of the SLC35F2/Qtp1 family, this protein is the sole protein responsible for q and Q import.

### Kinetic analysis of SLC35F2 q/Q transporter activity

Recent studies to determine the circulating levels of q in humans indicate a range of 1-10 nM, with higher levels reported in females (4, 25) but a surprising absence of detectable Q [4]. It therefore was of interest to determine the substrate affinity and specificity of SLC35F2, to provide an indication of its physiological activity. Of note, the SLC35F2 solute carrier has previously been identified as a unique, high-specificity transporter for the small molecule anti-cancer agent YM155 (sepantronium bromide), with retroviral disruption in KBM7 cells conferring drug resistance to this molecule (26).

To negate the interference of multiple downstream steps, the uptake kinetics of SLC35F2 for Q was determined in HeLa cells that were knocked out for QNG1 and therefore are incapable of converting Q to q (Fig. 4A, upper panel). Intracellular Q and Q-5′MP were quantified by LC-MS/MS and their additive concentration was used to measure uptake kinetics, providing a K_m_ of 174 nM and a Vmax of 0.87 pmol/min (lower panel). To examine q uptake, HeLa cells deficient in Qtrt1 were exploited (Fig. 4B, upper panel), providing a K_m_ of 67 nM for q uptake through both the SLC35F2 transporter and the unidentified secondary mechanism (lower panel). To obtain further data on the uptake kinetics of the unidentified transporter, we took advantage of the fact that YM155 is a high-specificity substrate for SLC35F2 uptake and thus capable of competitively inhibiting the transporter (Fig. 4C, upper panel) (26). Saturating concentrations of YM155 were added to the Qtrt1 deficient HeLa cells to inhibit q uptake, revealing that the second transporter for q has a higher K_m_ of 260 nM (lower panel), which agrees with previous literature wherein both a low-K_m_ and high-K_m_ transporter for the q nucleobase had been demonstrated (5). Together, the data indicate that the SLC35F2 transporter is uniquely tasked for Q uptake in humans with a K_m_ that is comparatively high relative to that of q and which could indicate differential roles for the transporter in the uptake of each form of the micronutrient.

**Figure 4.**
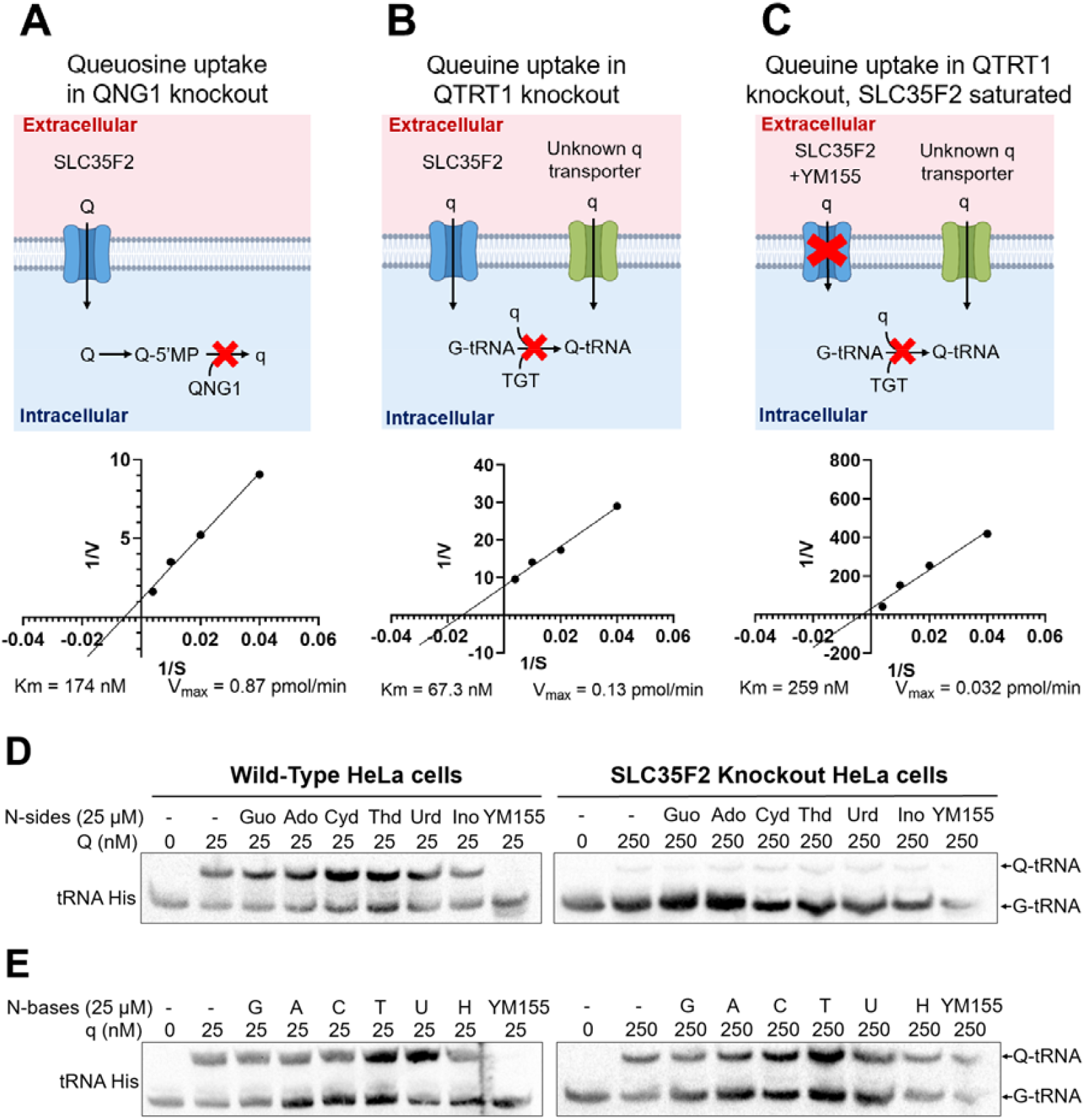
Analysis of q/Q uptake kinetics and specificity of SLC35F2 in human cells. **(A)** Uptake of Q was examined by adding increasing concentrations of Q (25, 50, 100, 250 nM) over four timepoints (15, 30, 45 and 60 mins) to QNG1-knockout HeLa cells, followed by methanol:water extraction from cell pellets and LC-MS/MS analysis to quantify the intracellular Q and Q-5′MP concentrations. (**B-C**) Similar analysis was performed for q in *Qtrt1*-knockout HeLa cells that were untreated (B) or administered the known SLC35F2 inhibitor YM155 (C) at saturating concentrations, before methanol:water extraction and LC-MS/MS analysis. **(D-E)** Import of q and Q by SLC35F2 was inhibited by YM155, but not other nucleobases or nucleosides. To wild-type or SLC35F2 knockout HeLa cells, a fixed concentration of Q (25 nM) or q (25 nM) was added, as indicated, in the absence (-) or presence of saturating concentrations of nucleobases or nucleosides (25 *μ*M) or YM155 (25 *μ*M). Total RNA was extracted and the Q-modification status of tRNA^His^ analysed by APB Northern blotting. Nucleobases - guanine (G), adenine (A), cytosine (C), thymine (T), uracil (U), hypoxanthine (H). Nucleosides - guanosine (Guo), adenosine (Ado), cytidine (Cyd), thymidine (Thy), uridine (Urd), inosine (Ino).

### The SLC35F2 transporter is highly specific for q/Q micronutrient uptake

Earlier studies showed that transporters for q are specific, and that uptake cannot be outcompeted by a variety of nucleobases or nucleotides (5). Using the level of Q-modified tRNA as an indirect indicator of the ability of cells to import Q or q, competitive uptake studies were performed using a range of nucleosides and nucleobases. In all cases in wild-type cells, none of the canonical nucleosides were capable of competing against Q-nucleoside uptake (Fig. 4D). Similarly, none of the tested nucleobases competed against q uptake (Fig. 4E) at saturating concentrations (25 μM versus 25 nM) except for the anti-cancer drug YM155, which effectively outcompeted Q/q import. This further validates SLC35F2 as the high-specificity transporter for both q and Q, both of which possess a cyclopentene-diol moiety not shared with canonical nucleobase or nucleoside molecules.

### Structural insights into SLC35F2/Qtp1 proteins

Human SLC35F2, *T. brucei* Qtp1 (ORF Tb927.4.320), and *S. pombe* Qtp1 (*SPCC320*.*08)* are all members of the SLC35F subfamily of the large solute carrier family 35 (SLC35). SLC35 proteins are highly conserved TM proteins typically composed of 10 α-helices (10-TM) arranged in two bundles of five helices each (12). These bundles are related by a pseudo-twofold symmetry axis running parallel to the membrane. Architecturally, they are part of the Drug/ Metabolite Exporter (DME) structural superfamily (27) and operate using an alternating-access mechanism to facilitate solute exchange across the membrane. This process involves two gates, formed by specific helical pairs, that alternately open and close on the opposite sides of the DME fold, enabling the central solute-binding site to access solutes from either side of the membrane. To date, the SLC35 family comprises 32 members classified into seven subfamilies (SLC35A-G) based on sequence similarity and phylogenetic analyses, with members within a subfamily differentiated by the solutes they transport. For the majority of SLC35 proteins (21 of the 32 members), the substrates are unknown (orphan transporters) [27], while most of the functionally characterized members (mostly in subfamilies A-D) are Nucleotide Sugar Transporters (NSTs) located in the Golgi and endoplasmic reticulum membranes, where they transport specific nucleotide sugars into the lumens of these organelles to serve as substrates for glycosylation reactions (28–31). NSTs use an antiport mechanism and typically utilize the corresponding luminal nucleoside monophosphates as their antiport molecules (32). Additionally, members of the SLC35B subfamily include an ATP/ADP exchanger (33) and 3′-phosphoadenosine 5′-phosphosulfate (PAPS) transporters (34). In the SLC35F subfamily, which comprises six members, none have been reported to function as an NST, and only one has been functionally characterized and found to transport thiamine across the plasma membrane (SLC35F3, (35)).

We built a structure-based multisequence alignment of human, *S. pombe* and *T. brucei* SLC35F2/Qtp1 proteins based on the superposition of tertiary structures predicted by the AI-based tool AlphaFold (36) (**Fig. S7**). Overall, the three proteins exhibit 14% sequence identity and 31% similarity. Their predicted structures share the common 10-TM transporter core and possess different, putatively extra-membranous, N- and C-terminal regions that could not be modeled with confidence. The 10-TM core is most similar to the crystal structure of the CMP-sialic acid transporter SLC35A1 (12, 37) (e.g., mouse CMP-sialic acid transporter, PDB ID 6OH3, r.m.s.d. 3.3 Å over 247 Cα atoms, 13% sequence identity (38)). (e.g., mouse CMP-sialic acid transporter, PDB ID 6OH3, r.m.s.d. 3.3 Å over 247 Cα atoms, 13% sequence identity (38)) In terms of their topology and orientation in the membrane, SLC35 proteins are classified as type III integral membrane proteins, i.e. their N- and C-termini face the cytoplasm (12). Notably, the predicted structures of human, *S. pombe* and *T. brucei* SLC35F2/Qtp1 proteins exhibit a biased distribution of positively charged amino acids (**Fig. S7B**), consistent with the “positive-inside rule” for integral membrane proteins, whereby basic amino acids cluster at the cytoplasmic interface (39). Analysis of the topology of the core membrane-spanning domain using the online tool Deep TMHMM (40) suggests that the N- and C-termini are located on the cytoplasmic side (**Fig. S7A**). These observations are consistent with the orientation of a type III integral membrane protein (**Fig. S7A**).

## Discussion

SLC35F2 belongs to the SLC35 solute superfamily and is a part of the F subfamily, which mostly consists of orphan transporters whose substrates and physiological function(s) are yet to be determined (30). Herein, we demonstrate by homozygous disruption in human HeLa cells that SLC35F2 functions as a key transporter for queuine and queuosine. We show that it is the sole high-affinity plasma membrane transporter for the Q nucleoside and the primary high-affinity transporter for the nucleobase q in human cells. Furthermore, it is the exclusive transporter of both q and Q in *T. brucei* and *S. pombe* and thus has been renamed Qtp1 in those two species.

Previous literature has shown that q uptake can occur by two different transporters in human cells, a low K_m_ (30 nM) component demonstrating rapid saturation and a high K_m_ component (350 nM) showing a slower linear uptake, but their identity was not determined (5). Here we demonstrate that SLC35F2 accounts for high affinity K_m_ q transporter activity (67 nM), and that there exists an independent lower affinity transporter for q (259 nM) that is evident from competitive inhibition of SLC35F2 with saturating concentrations of the anti-cancer compound YM155. Notably, the K_m_ for q is higher than the maximum levels determined in human serum under normal conditions of ∼25 nM (4, 25) and could therefore indicate that the levels of q in serum are capable of reaching higher levels, such as after a meal replete with q/Q. In the case of Q, its levels have not been detectable in human plasma (4). Therefore, the observed K_m_ of 174 nM could suggest SLC35F2’s role in Q uptake is limited to environments that are rich in this micronutrient, such as the human gut, where Q is liberated from tRNA turnover either from the microbiome or from food. In this regard, an analysis of tissue-specific expression data ((41) and Table S4) finds that there is a heavy propensity for SLC35F2 to be expressed in the human alimentary canal, suggesting that SLC35F2 is responsible for the active transport of Q/q during the digestive process. In our previous work, we observed that the liver has distinctly higher levels of the Q salvage enzyme QNG1 (18). It could therefore be envisaged that Q is transported from the gut by SLC35F2 and delivered to the liver for cleavage by QNG1, which then serves to liberate the q base for subsequent release into the serum and wider distribution throughout the body. Whether the SLC35F2 transporter also functions in a reversible manner to release q to the extracellular space is an outstanding question.

Similar to other eukaryotes, the blood borne parasite *T. brucei* is auxotrophic for q/Q, and it is interesting that it exploits a homologous transporter for uptake (termed Qtp1 in this work). In this regard, tRNA from blood-stream trypanosomes shows a greater level of Q-modification relative to the procyclic form that resides in the midgut of the tsetse fly (22), a difference that could potentially be explained by differential expression of the Qtp1 transporter.

The SLC35F2 gene was first identified during the construction of a transcription map of the ATM gene locus located on chromosome region 11q23, which is associated with the genetic disorder ataxia telangiectasia (42). Subsequently, an analysis of SLC35F2 mRNA expression by RT-PCR revealed that it is highest in human adult salivary glands (41). Previous studies on SLC35F2 have linked its expression to the progression of several forms of cancer (43). For example, high levels of SLC35F2 expression have been identified in non-small cell lung carcinoma tissue (under the name LSCC-3), suggesting its use as a prognostic biomarker (44). Moreover, elevated levels of SLC35F2 have been associated with papillary thyroid carcinoma (PTC) and inhibiting SLC35F2 expression was shown to suppress the malignant phenotype of PTC (45). Also, SLC35F2 shows oncogenic behavior, promoting cell proliferation and metastasis in H1299 lung cancer cells (46). Furthermore, SLC35F2 is highly expressed in human pluripotent stem cells, which underlies the high cytotoxicity of YM155 in these cells (47). Given our identification of SLC35F2 as an import molecule for q and Q, one interpretation of its oncogenic activity is that its overexpression increases cellular levels of q/Q and thus promotes higher Q34 modification of tRNAs. This in turn may give malignant cells a selective advantage over normal cells. Elucidating the role of SLC35F2 in q/Q import will deepen our understanding of its involvement in cancer progression and open new avenues for the development of targeted anti-cancer therapies.

## Materials and Methods

The *in silico* methods are given below. The growth conditions, mutant constructions, and analytical methods for Q/q detection and transport assays for HeLa, *T. brucei* and *S. pombe* cells are given in the supplemental methods section.

### Bioinformatic Analyses

The workflow for searching for candidates of Q precursor transmembrane transporters comprised three steps. First, QTRT^-^ and QTRT^+^ organisms were selected for comparative genomic analysis (Fig. S1 Step I). To identify the QTRT^-^ Eukaryotes, 245,896,766 protein sequences and metadata were retrieved from the Uniprot database (www.uniprot.org/help/downloads). Homologs of the human QTRT1 (Uniprot ID Q9BXR0) were searched from the proteomes of 10,768 taxa that contained at least 5000 protein entries using DIAMOND v2.1.8 (48) with the setting “--very-sensitive --matrix BLOSUM45 --evalue 0.001 -k0”. Four taxa of the Eukaryota superkingdom (including Sar, Viridiplantae, Fungi and Metazoa) that comprised at least ten QTRT^-^ organisms (Table S51) were subjected to further analyses. To ensure that all *Qtrt1* genes were called, *Qtrt1* coding sequences were retrieved using tBLASTn (49) searching the NCBI nt and wgs databases (50) limited to genomes from these four taxa. Organisms without hits at an E-value of 0.001 and 20% minimum identity were retained as QTRT^-^. Organisms encoding QTRT1 proteins between 300-500 amino acid in length were identified as QTRT^+^. These organisms were then sorted in descending order based on the number of proteins in any given taxa with at least four transmembrane domains, and this information was used to select approx. 200 organisms in each rank with criteria in Table S6. The QTRT^-^ and QTRT^+^ organisms are listed in Tables S3 and S7-9. Potential transporter families from both QTRT^+^ and QTRT-organisms were then clustered based on sequence similarity (using MMseqs2(51) (ref) with --cov-mode 0; Fig. S1 Step II). Clustering criteria for minimum coverage and sequence identity were tested in increments from 5 to 50%. A 30% threshold for both criteria was chosen for separation between homologous families as the number of proteins in each cluster sharply decreased at this cut-off (Table S10). Finally, the potential transporter subfamilies clusters were given scores as follows: the presence of a family member in a given QTRT^+^ organism or its absence in a given QTRT^-^ organism were scored by multiplying by 1; while the presence of a family member in a QTRT^-^ organism or its absence in a QTRT^+^ organism were penalized by multiplying by -1 (Fig. S1 Step III). The score of each family was normalized by the expected maximum score (present in all QTRT^+^ organisms and absent in QTRT^-^ organisms) so it fell within the range of -1 to 1. The scores of the top 200 candidates for the four ranks are listed in Table S1. The scripts used in the workflow is available at https://github.com/vdclab/published_scripts.

### Sequence Similarity Networks (SSNs)

SSNs were generated using the Enzyme Function Initiative (EFI) analytic suite (52) and visualized using Cytoscape (3.10.1) (53). 5782 sequences for the SLC35F1/F2 solute transporter family in Eukaryota were retrieved with the InterPro accession (54) of “IPR052221” through the “Family” option of EFI-EST (EFI Enzyme Similarity Tool)(52). The initial SSN was generated with an alignment score threshold (AST) set such that each connection (edge) represented a sequence identity above 40%. The obtained SSN was first colored by the presence and absence of QTRT in each organism. More stringent SSNs were then created by gradually increasing the AST in increments (usually by 10). This process was iterated until clusters containing nodes from QTRT^-^ organisms were separated from those comprising nodes from QTRT^+^ organisms. Clusters with less than 10 sequences were hidden for better visualization. The UniProt IDs of IPR05221 family sequences in the SSN are listed in Table S11.

### Phylogenetic Analysis of SLC35F1/F2 family Proteins

34 IPR05221 family sequences across different clades/kingdoms and three outgroup sequences, SLC35B1-4/HUT1 were aligned using MUSCLE v5.2 (55) and trimmed using BMGE v1.12 (56) with matrix BLOSUM30. The maximum likelihood tree was generated using FastTree v2.1.11 (57) with -lg -cat 20 -gamma model and bootstrap (1000 replicates) and visualized by iTOL v6.9.1 (58) The QTRT^-^ organisms are colored in red. The branches were colored by clades or kingdoms, and those with bootstrap support values higher than 0.7 were indicated by blue dots. The sequences used in the tree are listed in Table S12.

### Sequence and Structure Alignment

Structural models of SLC35F2 sequences from human, *S. pombe* strain 972 / ATCC 24843, and *T. brucei* strain 927/4 (Uniprot IDs Q8IXU6, O59785 and Q57UU3, respectively) were generated using AlphaFold v4 (36) and aligned in PROMALS3D (59). The resulting structure-based multi-sequence alignment was visualized using ESPript server v3.0 (60).

## Supporting information

SupplementalTablesS1toS13

## References

1. B. N. Ames, Prolonging healthy aging: Longevity vitamins and proteins. Proc Natl Acad Sci U S A 115, 10836–10844 (2018).

2. C. Fergus, D. Barnes, M. A. Alqasem, V. P. Kelly, The queuine micronutrient: Charting a course from microbe to man. Nutrients 7, 2897–2929 (2015).

3. J. R. Katze, Queuosine Metabolism: Possible relation to B-cell activation by C8 derivatives of guanosine. Proc Soc Exp Biol Med 179, 492–496 (1985).

4. X. Pan, et al., Development, validation and application of an LC–MS/MS method quantifying free forms of the micronutrients queuine and queuosine in human plasma using a surrogate matrix approach. Anal Bioanal Chem 416, 5711–5719 (2024).

5. M. S. Elliott, R. W. Trewyn, J. R. Katze, Inhibition of Queuine uptake in cultured human fibroblasts by phorbol-12,13-didecanoate1. Cancer Res 45, 1079–1085 (1985).

6. R. C. Morris, B. J. Brooks, K. L. Hart, M. S. Elliott, Modulation of queuine uptake and incorporation into tRNA by protein kinase C and protein phosphatase. Biochim Biophys Acta Mol Cell Res 1311, 124–132(1996).

7. M. S. Elliott, D. L. Crane, Protein kinase C modulation of queuine uptake in cultured human fibroblasts. Biochem Biophys Res Commun 171, 393–400 (1990).

8. R. Zallot, et al., Plant, animal, and fungal micronutrient queuosine is salvaged by members of the DUF2419 protein family. ACS Chem Biol 9, 1812–1825 (2014).

9. M. H. Saier, et al., The Transporter Classification Database (TCDB): 2021 update. Nucleic Acids Res 49, D461–D467 (2021).

10. M. Kanehisa, Y. Sato, M. Kawashima, M. Furumichi, M. Tanabe, KEGG as a reference resource for gene and protein annotation. Nucleic Acids Res 44, D457–D462 (2016).

11. A. Orellana, C. Moraga, M. Araya, A. Moreno, Overview of nucleotide sugar transporter gene family functions across multiple species. J Mol Biol 428, 3150–3165(2016).

12. S. Kamiyama, H. Sone, Solute Carrier Family 35 (SLC35)—An overview and recent progress. Biologics 4, 242–279 (2024).

13. R. Fredriksson, K. J. V. Nordström, O. Stephansson, M. G. A. Hägglund, H. B. Schiöth, The solute carrier (SLC) complement of the human genome: Phylogenetic classification reveals four major families. FEBS Lett 582, 3811–3816 (2008).

14. P. R. Bushel, et al., Dissecting the fission yeast regulatory network reveals phase-specific control elements of its cell cycle. BMC Syst Biol 3, 93 (2009).

15. F. Van den Bossche, et al., Residence of the Nucleotide Sugar Transporter Family Members SLC35F1 and SLC35F6 in the Endosomal/Lysosomal Pathway. Int J Mol Sci 25, 6718 (2024).

16. G. L. Igloi, H. Kossel, Affinity electrophoresis for monitoring terminal phosphorylation and the presence of queuosine in RNA. Application of polyacrylamide containing a covalently bound boronic acid. Nucleic Acids Res 13, 6881–6898 (1985).

17. M. Müller, et al., Queuine links translational control in eukaryotes to a micronutrient from bacteria. Nucleic Acids Res 47, 3711–3727 (2019).

18. S. H. Hung, et al., Structural basis of Qng1-mediated salvage of the micronutrient queuine from queuosine-5′-monophosphate as the biological substrate. Nucleic Acids Res 51, 935–951 (2023).

19. B. I. Patel, M. Heiss, A. Samel-Pommerencke, T. Carell, A. E. Ehrenhofer-Murray, Queuosine salvage in fission yeast by Qng1-mediated hydrolysis to queuine. Biochem Biophys Res Commun 624, 146–150 (2022).

20. A. E. Ehrenhofer-Murray, Cross-Talk between Dnmt2-dependent tRNA methylation and queuosine modification. Biomolecules 7, 14 (2017).

21. S. Kulkarni, et al., Preferential import of queuosine-modified tRNAs into Trypanosoma brucei mitochondrion is critical for organellar protein synthesis. Nucleic Acids Res 49, 8247–8260 (2021).

22. S. Dixit, et al., Dynamic queuosine changes in tRNA couple nutrient levels to codon choice in Trypanosoma brucei. Nucleic Acids Res 49, 12986–12999 (2021).

23. A. C. Kessler, et al., Retrograde nuclear transport from the cytoplasm is required for tRNATyr maturation in T. brucei. RNA Biol 15, 528–536 (2018).

24. K. M. Rutherford, M. Lera-Ramírez, V. Wood, PomBase: a Global Core Biodata Resource—growth, collaboration, and sustainability. Genetics 227, iyae007 (2024).

25. P. Richard, et al., Queuine, a bacterial-derived hypermodified nucleobase, shows protection in in vitro models of neurodegeneration. PLoS One 16, e0253216 (2021).

26. G. E. Winter, et al., The solute carrier SLC35F2 enables YM155-mediated DNA damage toxicity. Nat Chem Biol 10, 768–773 (2014).

27. P. M. Berninsone, C. B. Hirschberg, Nucleotide sugar transporters of the Golgi apparatus. Curr Opin Struct Biol 10, 542–547 (2000).

28. K. T. Schjoldager, Y. Narimatsu, H. J. Joshi, H. Clausen, Global view of human protein glycosylation pathways and functions. Nat Rev Mol Cell Biol 21, 729–749(2020).

29. Z. Song, Roles of the nucleotide sugar transporters (SLC35 family) in health and disease. Mol Aspects Med 34, 590–600(2013).

30. B. Hadley, et al., Nucleotide Sugar Transporter SLC35 family structure and function. Comput Struct Biotechnol J 17, 1123–1134(2019).

31. S. Mikkola, Nucleotide Sugars in Chemistry and Biology. Molecules 25, 5755 (2020).

32. D. L. Jack, N. M. Yang, M. H. Saier, The drug/metabolite transporter superfamily. Eur J Biochem 268, 3620–3639 (2001).

33. M.-C. Klein, et al., AXER is an ATP/ADP exchanger in the membrane of the endoplasmic reticulum. Nat Commun 9, 3489 (2018).

34. C. B. Hirschberg, P. W. Robbins, C. Abeijon, Transporters of nucleotide sugars, ATP, and nucleotide sulfate in the endoplasmic reticulum and Golgi apparatus. Annu Rev Biochem 67, 49–69 (1998).

35. K. Zhang, et al., Genetic implication of a novel thiamine transporter in human hypertension. J Am Coll Cardiol 63, 1542–1555 (2014).

36. J. Jumper, et al., Highly accurate protein structure prediction with AlphaFold. Nature 596, 583–589 (2021).

37. E. Nji, A. Gulati, A. A. Qureshi, M. Coincon, D. Drew, Structural basis for the delivery of activated sialic acid into Golgi for sialyation. Nat Struct Mol Biol 26, 415–423 (2019).

38. S. Ahuja, M. R. Whorton, Structural basis for mammalian nucleotide sugar transport. Elife 8, e45221 (2019).

39. C. Baeza-Delgado, M. A. Marti-Renom, I. Mingarro, Structure-based statistical analysis of transmembrane helices. Eur Biophys J 42, 199–207 (2013).

40. J. Hallgren, et al., DeepTMHMM predicts alpha and beta transmembrane proteins using deep neural networks. bioRxiv 2022.04.08.487609 (2022).

41. M. Nishimura, S. Suzuki, T. Satoh, S. Naito, Tissue-Specific mRNA Expression Profiles of Human Solute Carrier 35 Transporters. Drug Metab Pharmacokinet 24, 91–99 (2009).

42. T. Stankovic, et al., Construction of a transcription map around the gene for ataxia telangiectasia: identification of at least four novel genes. Genomics 40, 267–276 (1997).

43. R. Kotolloshi, M. Hölzer, M. Gajda, M. O. Grimm, D. Steinbach, SLC35F2, a transporter sporadically mutated in the untranslated region, promotes growth, migration, and invasion of bladder cancer cells. Cells 10, 1–12 (2021).

44. L. Bu, G. Jiang, F. Yang, J. Liu, J. Wang, Highly expressed SLC35F2 in non-small cell lung cancer is associated with pathological staging. Mol Med Rep 4, 1289–1293 (2011).

45. J. He, et al., Solute carrier family 35 member F2 is indispensable for papillary thyroid carcinoma progression through activation of transforming growth factor-β type I receptor/apoptosis signal-regulating kinase 1/mitogen-activated protein kinase signaling axis. Cancer Sci 109, 642–655 (2018).

46. X. Li, et al., Influence on the behavior of lung cancer H1299 cells by silencing SLC35F2 expression. Cancer Cell Int 13, 1–6 (2013).

47. K.-T. Kim, et al., Safe scarless cassette-free selection of genome-edited human pluripotent stem cells using temporary drug resistance. Biomaterials 262, 120295 (2020).

48. B. Buchfink, K. Reuter, H. G. Drost, Sensitive protein alignments at tree-of-life scale using DIAMOND. Nat Methods 18, 366–368 (2021).

49. S. F. Altschul, W. Gish, W. Miller, E. W. Myers, D. J. Lipman, Basic local alignment search tool. J Mol Biol 215, 403–410 (1990).

50. E. W. Sayers, et al., Database resources of the national center for biotechnology information. Nucleic Acids Res 50, D20–D26 (2022).

51. M. Steinegger, J. Söding, MMseqs2 enables sensitive protein sequence searching for the analysis of massive data sets. Nat Biotechnol \35, 1026–1028(2017).

52. R. Zallot, N. Oberg, J. A. Gerlt, The EFI web resource for genomic enzymology tools: leveraging protein, genome, and metagenome databases to discover novel enzymes and metabolic pathways. Biochemistry 58, 4169–4182 (2019).

53. P. Shannon, et al., Cytoscape: A software environment for integrated models of biomolecular interaction networks. Genome Res 13, 2498–2504 (2003).

54. T. Paysan-Lafosse, et al., InterPro in 2022. Nucleic Acids Res 51, D418–D427 (2023).

55. R. C. Edgar, Muscle5: High-accuracy alignment ensembles enable unbiased assessments of sequence homology and phylogeny. Nature Commun 13, 1–9 (2022).

56. A. Criscuolo, S. Gribaldo, BMGE (Block Mapping and Gathering with Entropy): A new software for selection of phylogenetic informative regions from multiple sequence alignments. BMC Evol Biol 10, 1–21 (2010).

57. M. N. Price, P. S. Dehal, A. P. Arkin, FastTree 2 – Approximately maximum-likelihood trees for large alignments. PLoS One 5, e9490 (2010).

58. I. Letunic, P. Bork, Interactive Tree of Life (iTOL) v6: Recent updates to the phylogenetic tree display and annotation tool. Nucleic Acids Res 52, W78–W82 (2024).

59. J. Pei, M. Tang, N. V. Grishin, PROMALS3D web server for accurate multiple protein sequence and structure alignments. Nucleic Acids Res 36, W30–W34 (2008).

60. X. Robert, P. Gouet, Deciphering key features in protein structures with the new ENDscript server. Nucleic Acids Res 42, W320–W324 (2014).

61. X. Zhao, et al., Glycosylated queuosines in tRNAs optimize translational rate and post-embryonic growth. Cell 186, 5517–5535.e24 (2023).

