## SupplementalsMethodsandFigures for "The oncogene SLC35F2 is a high-specificity transporter for the micronutrients queuine and queuosine"

[&]Current address: eSTEAMed Learning Inc., Maitland, FL 32751, USA

**Supplementary tables**

**Table S1.** The list of candidates in each of the four taxonomic ranks.

**Table S2.** The *S. pombe* Qtp1 encoding gene *SPCC320.08* is co-expressed with translation genes.

**Table S3**. Viridiplantae used for comparative genomic searching for Q transporter.

**Table S4.** Expression levels of SLC35F2 in human tissues.

**Table S5.** Searching of QTRT- and QTRT+ organisms in Uniprot database in selected taxonomic ranks.

**Table S6.** Criteria used to select QTRT+ organisms for each of the four ranks.

**Table S7**. Fungi used for comparative genomic searching for Q transporter.

**Table S8**. Metazoa used for comparative genomic searching for Q transporter.

**Table S9**. Sar used for comparative genomic searching for Q transporter.

**Table S10.** The average size of clusters when applying different clustering criteria using mmseqs2.

**Table S11.** Sequences used in the Sequence Similarity Network (SSN).

**Table S12.** Sequences used in the phylogenetic tree of 34SLC35F Transporter from the InterPro IPR052221 family.

**Table S13.** Summary of Multiple Reaction Monitoring (MRM) transition settings for Q metabolites.

### Supplemental Methods

**HeLa cell culture.** Cells were purchased from the European Collection of Cell Cultures (ECACC), grown in an atmosphere of 5% CO2 at 37 $^{o}$C and routinely maintained in Dulbecco's Modified Eagle's Medium (DMEM) media containing 10% fetal bovine serum (Sigma), 1.5 mM L -glutamine (Gibco), 1% penicillin/streptomycin (Gibco), except for studies evaluating q/Q salvage, where cells were grown in OptiPro TM Serum-free media (Gibco) supplemented with 2 mM L-glutamine for three or more passages before experiments. Chemically synthesized q was a gift from Dr. Susumu Nishimura (Tsukuba University, Japan) or purchased from Epitoire (Singapore). Queuosine was purchased from Epitoire (Singapore).

**Preparation of HeLa cell and media extracts for LC-MS/MS analysis.** Cells were grown on 6-well plates in 2 ml of OptiPro serum-free media containing varying concentrations of q/Q and nucleobases, nucleosides or YM155, as indicated. Media extracts were prepared by adding 400 μl methanol to 100 μl of growth media, vortexing, centrifuging at 5,800 x g, 4 °C for 5 minutes and collecting the supernatants. Pellets from the media samples were extracted twice more with 100 μl ice-cold MeOH:$H_2O$ (80:20 v/v) and the combined extracts were dried down using a Thermo Scientific Savant SpeedVac Vacuum Concentrator at 2 mbar for 2 hours at 40 °C. For preparation of cytosolic extracts, cells were washed twice with 1 ml ice-cold phosphate-buffered saline, and 500 μl of ice-cold MeOH-$H_2O$ (80:20 v/v) was added directly onto the cells, which were scraped and transferred to an Eppendorf tube. The cells were vortexed, and the homogenate flash frozen in liquid nitrogen for 2 min followed by thawing in an ice bath for 10 min. The sample was centrifuged at 5,800 x g for 5 min at 4 °C and the supernatant was collected. The pellets arising from the cytosol samples were extracted twice more with 250 μl ice-cold MeOH-$H_2O$ (80:20 v/v) and the combined extracts were SpeedVac-dried as above. All extracts were reconstituted in 100 μl water for LC-MS/MS analysis. To account for the total volume of media used in cell culture (2 ml), analyte quantities measured in the 100-μl media extracts were multiplied by 20.

**LC-MS/MS quantification of free q, Q and Q-5′MP in HeLa cell extracts and media.** Stock solutions of queuine and queuosine were made to a concentration of 1 mM and Q-5′MP was prepared at the concentration of 3 mM, all in water and stored at -20 °C until use. The calibration and quality control (QC) solutions were prepared freshly in ultrapure water on the day of analysis. Eight calibration standards with a concentration 1 – 0.0003 µM for queuine and queuosine; and 3 – 0.001 µM for Q-5′MP were prepared by serial dilution. QC solutions contained queuine and queuosine at 0.8, 0.02 and 0.0008 µM; and Q-5′MP at 2.4, 0.06 and 0.0024 µM. The resulting standard curves were used for quantification of Q, q (Fig. S8) and Q-5'MP (Fig. S9) in experimental samples. HeLa cell and media extracts, reconstituted in 100 µl water, were analyzed using a LC-MS/MS system consisting of an AB SCIEX ExionLC system (Foster, California, USA) coupled to an AB SCIEX Triple Quad 5500+ mass spectrometer (Foster, California, USA). The mass spectrometer was operated in ESI positive mode using Multiple Reaction Monitoring (MRM). Analyte separation was performed on a Waters XSelect HSS T3 column (100 × 4.6 mm, 3.5 µm, Milford, MA, USA) at 45 °C using a gradient of 0.1% formic acid in water (mobile phase A) and 0.1% formic acid in acetonitrile (mobile phase B) at a flow rate of 1.0 ml/min. The mobile gradient was set as follows: 2% B to 3% B from 0 to 3 min, ramped to 98% B at 4 min and hold for 2 min, decrease from 98% to 2% B in 0.5 min and then equilibrated for 3.5 min at 2% B. The injection volume was 5 µl. Mass spectrometric parameters setting were as follows: curtain gas 30; collision gas 9; ion spray voltage 5500; temperature 550; ion source gas 1, 50; ion source gas 2, 50; and entrance potential 10. The MRM transition settings are summarized in Table S13**.**

**CRISPR targeting of SLC35F2 in HeLa cells**

The SLC35F2 knock-in template was amplified by PCR from the p2attPC plasmid (Addgene ID #51547) using the forward primer 5′-AGATGTTGTCCTTGTGTATATGTGGGACAGCCATCGGCTCTGGCGGCGGAAGCGGAATGGCTACCGAGTACAAGCCCACG -3′ and the reverse primer 5′-ATGGGGGTGTTCACTTTGTATCTTTCTGCCAAATACCATAGAGCCCACCGCATCCCCAG-3′ according to a thermocycle of 98 °C for 30 s, followed by 98 °C for 10 s, 72 °C for 30s for 35 cycles. Homology arms (underlined solid line) and a spacer (highlighted in grey) were introduced as overhangs into the repair-template. PCR products were purified using the QIAquick gel PCR kit (Qiagen). A CRISPR-Cas9 targeting site was identified in the SLC35F2 gene (5′-TCTTTCTGCCAAATACTGGC-3′) using the CRISPOR online tool (http://crispor.tefor.net/). Equimolar concentrations (12 µM) of recombinant Cas9 protein from S. pyogenes (PNA Bio Ltd.) was annealed to tracrRNA and crRNA (Sigma) and delivered together with linear dsDNA repair template (3.6 µg) to HeLa cells ($6 \times 10^5$) using the Neon™ Transfection System (Thermo Fisher Scientific); 1005 V, 35 ms, 2 pulses. Electroporated cells were transferred to a 10 cm dish containing 10 ml of complete media and allowed to grow to ~90% confluency before treatment with puromycin (3 µg/ml) for 72 h. Cells were washed with PBS twice and resistant colonies allowed to grow for a further 4–5 days. Colonies were harvested using 8 mm cloning cylinders (C3983, Sigma) and expanded for analysis.

**APB-Northern assay for Q-tRNA detection in HeLa cells**

The queuosylation status of tRNA was determined using 3-AcrylamidoPhenylBoronic acid (APB; TCI chemicals Ltd) gels as previously described.(1) Briefly, following incubation of HeLa cells with synthetic q or Q for 24 h, RNA was extracted using the mirVana miRNA Isolation Kit (Ambion, AM1561), deacylated in 100 mM Tris-HCl (pH 9.0), added to 2x gel loading buffer II (AM8546G) and denatured at 95 °C for 5 min, before placing on ice. RNA was separated on a 30% APB-PAGE gel, run in 1x TAE buffer for 30 min at 250 V. The RNA was transferred to an Amersham Hybond membrane (GE Healthcare) at 15 V for 90 min in a cold room using a semi-dry transfer cell, the membrane rinsed in 1x TAE buffer and crosslinked to the membrane at 120,000 µJ (UV Stratalinker 1800). Membranes were pre-hybridised in ULTRAhyb buffer (AM8670, Ambion) at room temperature for 30 min. Locked Nucleic Acid (LNA) probes (Qiagen) for human tRNA$^{His}$ (5′-TGCGGCCACAACGCAGAGTACTAA-3′) 3′-OH-modified with DIG-11-dUTP and dATP using the DIG Oligonucleotide Tailing Kit (Roche). LNA probe was denatured at 95 ºC for 1 min and added to the membrane in ULTRAhyb buffer (5 nM final) and hybridized overnight at room temperature. Following hybridization, the membrane was washed twice with Low Stringent Buffer (2x SSC, 0.1% (w/v) SDS) for 15 min each, twice with High Stringent Buffer (0.1x SSC, 0.1% (w/v) SDS) for 15 min each and rinsed with Washing Buffer (1x SSC) for 5 min. The membrane was incubated in a Blocking Buffer for 3 h at room temperature, before addition of anti-DIG alkaline phosphatase-conjugated antibody (Roche, 11093274910, 1:15000 dilution) and incubation at room temperature for 45 min. The membrane was washed in DIG Washing Buffer four times (15 min each), incubated in Development Buffer for 5 min, and then incubated in 1 ml of CSPD substrate solution for 5 min. Photoemissions from dephosphorylation of CSPD (Applied Biosystems, T2217) (maximum intensity 475 nm) were detected using a Bio-Rad ChemiDoc Imaging system and Image Lab Software. Washing, blocking and detection buffers are all a part of the DIG Wash and Block Buffer Set (Roche, 11585762001).

**Genotyping screen of the SLC35F2 candidate knockout clones**

Genomic DNA was isolated from cells using the Gentra Puregene Cell Kit (Qiagen) following manufacturer's instructions. Genotyping analysis of SLC35F2 knock-out clones was completed by PCR amplification of the targeted alleles from isolated genomic DNA using the GoTaq PCR Core System kit (Promega, M7660) according to the manufacturer's guidelines. To test for the SLC35F1 WT allele the forward (5'-GATGCACCCGGCTGCGTTCTGATC-3') and reverse (5'-CAGCACTTTGCGGATCCTCTGG-3') primers were used for amplification. To test for the SLC35F2 WT allele the forward (5'-CATGAAATGAAGAACCACC TGG-3') and reverse (5'-CCTGATCGAAATGCCAGCATCAC-3') primers were used for amplification. All primers used have an annealing temperature of 60 °C.

***T. brucei* cell culturing and gene deletions**

*T. brucei* SmOx-Cas9 procyclic cells (2) were cultured at 27°C in SDM-79 media (3) supplemented with 10% Fetal Bovine Serum (FBS), and growth was assessed by counting with Neubauer chambers. Gene deletion was performed using a pPOTv4 plasmid protocol (4). Briefly, PCR-generated DNA cassettes targeting upstream and downstream regions in the Qtp1 ORF were electro-transfected into *T. brucei* cells alongside PCR-generated cassettes corresponding to hygromycin and neomycin resistance genes. Gene deletion was induced by the addition of 1 µg/mL tetracycline to the media overnight, generating sgRNAs that guided Cas9-produced DNA double-strand breaks, allowing the replacement of Qtp1 targeted loci with drug cassettes via UTR homologous recombination. Transformants were then selected with the addition of hygromycin and neomycin to 50 and 12 µg/ml to the media respectively. The deletion confirmation was performed by PCR amplification of WT and drug cassette ORFs. Oligos utilized for the pPOTv4 plasmid protocol were generated as follows: UpstreamsgRNAFor (5'-GAAATTAATACGACTCACTATAGGAAGGGAGGGCAATTTTCAGGGTTTTAGAGCTAGAAA TAGC-3'); DownstreamsgRNAFor (5'-GAAATTAATACGACTCACTATAGGAGACGGTAAACGTCTCCCCGGTTTTAGAGCTAGAAA TAGC-3'); sgRNARev (5'-AAAAGCACCGACTCGGTGCCACTTTTTCAAGTTGATAACGGACTAGCCTTATTTTAACTTG CTATTTCTAGCTCTAAAAC-3'); NeomycinFor (5'-ACCTACGAGTTAGTAACACCTAAAGCAGGGGTATAATGCAGACCTGCTGC-3'); NeomycinRev

(5’-CTCCCCGGCGACTTCCGTTGCGGTCAACCGCCGGAACCACTACCAGAACC-3’); HygromycinFor (5’-ACCTACGAGTTAGTAACACCTAAAGCAGGGGGTTCTGGTAGTGGTTCC-3’); HygromycinRev (5’-CTCCCCGGCGACTTCCGTTGCGGTCAACCGCCAATTTGAGAGACCTGTGC-3’).

#### Depletion of Q-tRNA and nutrient supplementation in *T. brucei*

Depletion of Q-tRNA$^{Tyr}$ was achieved by growing cells in media supplemented with 10% dialyzed FBS for 10 days. Absence of Q-tRNA$^{Tyr}$ was confirmed by APB-Northern blot hybridization performed on extracted total RNA samples revealing both Q-tRNA$^{Tyr}$ and G-tRNA$^{Tyr}$. Nutrient supplementation to rescue Q-tRNA was performed by addition of either queuosine or queuine to the media 24 hours prior to RNA isolation.

#### APB-Northern assay for Q-tRNA detection in *T. brucei*

Prior to gel electrophoresis, the Isolated RNA was deacylated by incubating in 0.1 M Tris-HCl pH 9.0 for 30 min at 37$^0$C. The APB gels were prepared by supplementing 50 mg of 3-aminophenylboronic acid (Sigma) per 10 ml of 8% acrylamide solution with 8M Urea in 1x TAE buffer. Then, 5 $\mu$g of RNA of each sample was resolved on APB gels at 75 V for 5.5 hrs. Prior to the transfer, they were stained with ethidium bromide for 10 min and visualized. Then, RNA resolved on gels were transferred into Zeta probe nylon membranes at 80 V for 2 hrs. The RNA was crosslinked into resulting blots by exposing to UV radiation for 1 min. Northern blot hybridization was carried out according to the manufacturer’s protocol (Bio-Rad). The northern blot probe for tRNA$^{Tyr}$ (5’-CCTTCCGGCCGGAATCGAACCAGCGAC-3’) was end-labeled with P$^{32}$ and was utilized for hybridization.

#### *S. pombe* strains and growth conditions

The *S. pombe* strains used in this study are derived from AEP580 (FY7385, h$^-$ leu1-32 ura4-D18 his3-D3). The *qtp1*$^+$ (SPCC320.08) gene deletion was obtained by homologous recombination-mediated replacement of the open reading frame with NatMX (5). The deletion strain was named AEP622 (*h*$^-$ *leu1-32 ura4-D18 his3-D3 qtp1*$\Delta$*::NatMX).* Cells were grown in YES medium (5 g/l yeast extract, 30 g/l glucose, 250 mg/l adenine, 250 mg/l histidine, 250 mg/l leucine, 250 mg/l uracil, 250 mg/l lysine) at 30 °C and supplemented with the synthetic nucleobase queuine (10 nM or 100 nM) or the synthetic nucleoside queuosine (150 nM) as indicated. To amplify the 5’ homology region of *qtp1* for generating the deletion cassette: (5’-GTGTTTACAAGTTTAGTATCTTGATTCGTTTTCTACG-3’) and Reverse (5’-GGGTATTCTGGGCCTCCATGTCGGAAGGCCAATATCATGTTCAAACGG-3’). To amplify NatMX with flanks homologous to 5’ and 3’ regions of *qtp1*$^+$:
Forward (5’--CCGTTTGAACATGATATTGGCCTTCCGACATGGAGGCCCAGAATACCC-3’) and Reverse (5’CCAAGCTCCAAAAATCGGAAGTCCAGTATAGCGACCAGCATTCACATACG--3’).
To amplify the 3’ homology region of *qtp1* for generating the deletion cassette: Forward (5’-CGTATGTGAATGCTGGTCGCTATACTGGACTTCCGATTTTTGGAGCTTGG-3’) and Reverse (5’-CGACAAAAATCGTAAGAGGTTCGTTTTTAACAC-3’). To verify the correct integration of the *qtp1*$\Delta$*::NatMX* deletion cassette from 5’-end: Forward (5'-CACGTACTGTATGTCTGATAACAAGTGG-3') and Reverse (5'-GCTAAATGTACGGGCGACAGTC-3’). To verify the correct integration of the *qtp1*$\Delta$*::NatMX* deletion cassette from 3’-end: Forward (5'-GACATCATCTGCCCAGATGCG-3') and Reverse (5'-CAATAATAACATACTGTAGGTCCTTTACCAGAAACG-3').

**APB-Northern assay for Q-tRNA detection in *S. pombe***

APB Northern blotting was performed as previously described (6, 7) with some modifications. Briefly, total RNA was isolated from 10 OD equivalents of cells grown in exponential phase using acidic phenol-chloroform extraction. 500 ng of RNA were deacylated by incubation in 10 mM Tris (pH 9) at 37 °C for 30 minutes. The deacylated RNA was supplemented with 1x RNA loading dye (Thermo Scientific) and denatured at 70 °C for 5 minutes. The samples were loaded on 12% urea gels containing 0.5 % APB. The gel was run for 2 hours at 30 mA in 1x TBE buffer. RNA was then transferred to positively charged Biodyne B 0.45 mm Nylon membrane (Pall Corporation) in 0.5x TBE at 150 mA for 90 minutes at 4 °C by wet transfer. RNA was fixed on the membrane by baking the membrane at 60 °C for 30 minutes and crosslinking with 120 $mJ/cm^2$ UV. $tRNA^{Asp}$ was detected using the North2South Chemiluminescent Hybridization and Detection Kit (Thermo Scientific). The initial membrane blocking was performed with DIG Easy Hyb (Roche). Hybridization was performed at 60ºC with a biotin-labeled probe for *S. pombe* $tRNA^{Asp}$ (5'-3' orientation) (Biotin-GGGCTGCAAGCGTGACAGG) at 50 ng/ ml. The blot was exposed using the ChemiDoc Imaging system (BioRad).

**References**


1. S. H. Hung, *et al.*, Structural basis of Qng1-mediated salvage of the micronutrient queuine from queuosine-5′-monophosphate as the biological substrate. *Nucleic Acids Res* **51**, 935–951 (2023).

2. T. Beneke, *et al.*, A CRISPR Cas9 high-throughput genome editing toolkit for kinetoplastids. *R Soc Open Sci* **4**, 170095 (2017).

3. R. Brun, Schönenberger, Cultivation and in vitro cloning or procyclic culture forms of *Trypanosoma brucei* in a semi-defined medium. Short communication. *Acta Trop* **36**, 289–92 (1979).

4. S. Dean, *et al.*, A toolkit enabling efficient, scalable and reproducible gene tagging in trypanosomatids. *Open Biol* **5**, 140197 (2015).

5. A. Wach, A. Brachat, R. Pöhlmann, P. Philippsen, New heterologous modules for classical or PCR-based gene disruptions in *Saccharomyces cerevisiae*. *Yeast* **10**, 1793–1808 (1994).

6. G. L. Igloi, H. Kössel, Affinity electrophoresis for monitoring terminal phosphorylation and the presence of queuosine in RNA. Application of polyacrylamide containing a covalently bound boronic acid. *Nucleic Acids Res* **13**, 6881–6898 (1985).

7. B. I. Patel, M. Heiss, A. Samel-Pommerencke, T. Carell, A. E. Ehrenhofer-Murray, Queuosine salvage in fission yeast by Qng1-mediated hydrolysis to queuine. *Biochem Biophys Res Commun* **624**, 146–150 (2022).

### Supplementary figures

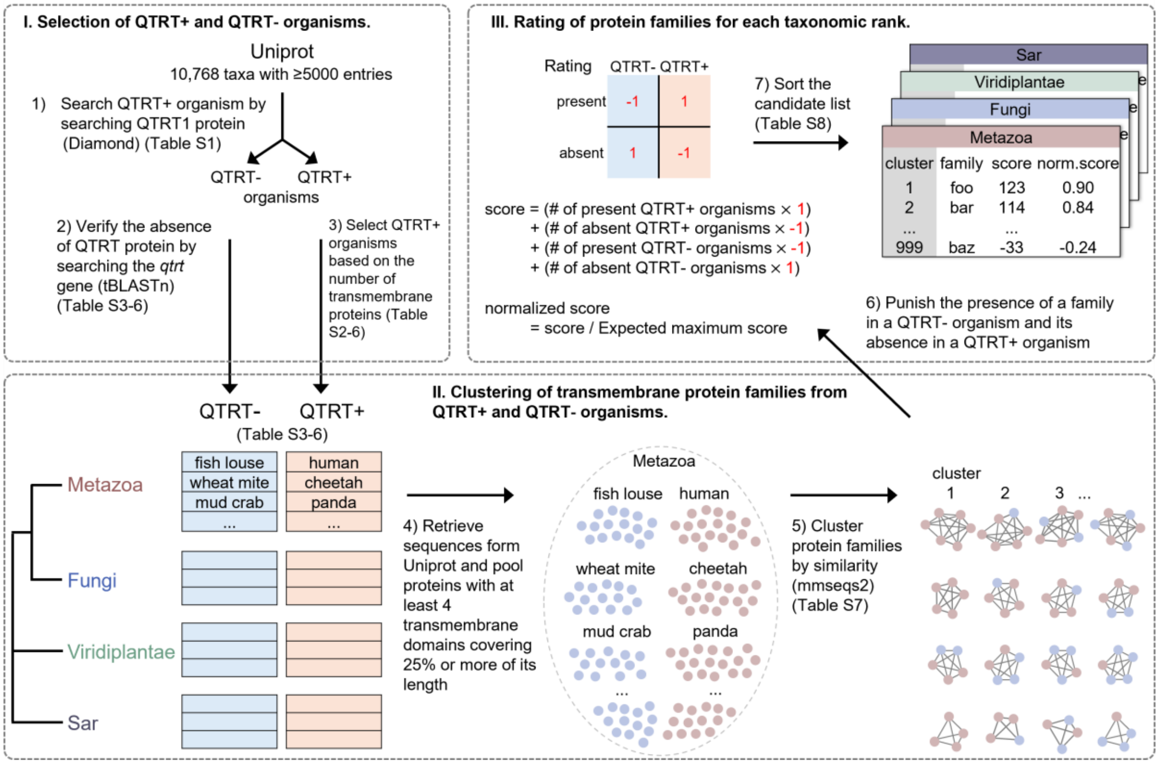


**Figure S1. The workflow for identifying Q precursor transporters in Eukaryota.** Tools used in each step and corresponding supplementary table are in parentheses. Steps 4-7 are repeated for each of the four selected taxonomic ranks, using Metazoa as an illustrative example. Details are provided in the Methods section.

**A**

Summary of expression profile of SLC35F1 (source: https://www.proteinatlas.org/)

Tissue, cell type and cancer cell line expression profile

| | |
|---|---|
| Tissue specificity | Group enriched (Brain, Retina) |
| Tissue expression cluster | Neuronal - Synaptic function (mainly) |
| Brain specificity | Low human brain regional specificity |
| Brain expression cluster | Neurons - Mixed function (mainly) |
| Single cell type specificity | Group enriched (Oligodendrocyte precursor cells, Inhibitory neurons, Excitatory neurons, Astrocytes) |
| Single cell type expression cluster | Oligodendrocyte precursor - Unknown function (mainly) |
| Tissue cell type classification | Cell type enriched (Colon - Enteric glia cells, Spleen - Fibroblast_1) |
| Immune cell specificity | Not detected in immune cells |
| Immune cell expression cluster | Not detected - no cluster assigned |
| Prognostic summary | SLC35F1 is a prognostic marker in Kidney renal clear cell carcinoma |
| Cancer specificity | Group enriched (Glioblastoma Multiforme, Skin Cuteneous Melanoma) |
| Cell line expression cluster | Skin cancer - ECM organization (mainly) |
| Cell line specificity | Cancer enhanced (Skin cancer) |

Top 15 nearest neighbours based on tissue RNA expression

| Neighbour | Description | Correlation | Cluster |
|---|---|---|---|
| APC | APC regulator of WNT signaling pathway | 0.9596 | 17 |
| LRRC4C | Leucine rich repeat containing 4C | 0.9404 | 17 |
| NEFL | Neurofilament light chain | 0.9404 | 17 |
| CACNG3 | Calcium voltage-gated channel auxiliary subunit gamma 3 | 0.9351 | 17 |
| SLC1A3 | Solute carrier family 1 member 3 | 0.9316 | 17 |
| DRD1 | Dopamine receptor D1 | 0.9298 | 17 |
| DCLK2 | Doublecortin like kinase 2 | 0.9246 | 33 |
| GRIA3 | Glutamate ionotropic receptor AMPA type subunit 3 | 0.9211 | 17 |
| VSTM2A | V-set and transmembrane domain containing 2A | 0.9193 | 17 |
| ELFN2 | Extracellular leucine rich repeat and fibronectin type III domain containing 2 | 0.9175 | 33 |
| FEZF2 | FEZ family zinc finger 2 | 0.9158 | 33 |
| PCDH11Y | Protocadherin 11 Y-linked | 0.9140 | 17 |
| C1orf216 | Chromosome 1 open reading frame 216 | 0.9140 | 17 |
| LHX2 | LIM homeobox 2 | 0.9123 | 17 |
| TNR | Tenascin R | 0.8930 | 33 |

Cluster 17: Neuronal - Synaptic function
Cluster 33: Brain - Neuronal signaling

Summary of SLC35F1 co-expressing genes (source: https://seek.princeton.edu/seek/)

Top 10 SLC35F1 co-expressing genes in top weighted datasets

| Rank | Z-score of Pearson correlation | Gene |
|---|---|---|
| 1 | 2.846 | NRXN1 |
| 2 | 2.560 | ZNF536 |
| 3 | 2.538 | GNAO1 |
| 4 | 2.509 | PPFIA2 |
| 5 | 2.499 | TRIM9 |
| 6 | 2.478 | GPM6B |
| 7 | 2.454 | ELAVL3 |
| 8 | 2.451 | POU3F2 |
| 9 | 2.447 | AMPH |
| 10 | 2.445 | NLGN1 |

Gene set enrichment analysis (GO (BP) term) of top 100 SLC35F1 co-expressing genes

| Term | p-value ? | q-value ? |
|---|---|---|
| multicellular organismal signaling | 5.68E-4 | 4.28E-3 |
| synaptic transmission | 4.18E-3 | 3.1E-2 |
| generation of neurons | 2.19E-2 | 1.56E-1 |
| ion transmembrane transport | 2.42E-2 | 1.56E-1 |
| axonogenesis | 2.42E-2 | 1.67E-1 |

**B**

Summary of expression profile of SLC35F2 (source: https://www.proteinatlas.org/)

Tissue, cell type and cancer cell line expression profile

| | |
|---|---|
| Tissue specificity | Tissue enhanced (Salivary gland) |
| Tissue expression cluster | Stomach - Proteolysis (mainly) |
| Brain specificity | Low human brain regional specificity |
| Brain expression cluster | Astrocytes - Mixed function (mainly) |
| Single cell type specificity | Cell type enhanced (Serous glandular cells, Basal keratinocytes, Suprabasal keratinocytes) |
| Single cell type expression cluster | Serous glandular cells - Salivary secretion (mainly) |
| Tissue cell type classification | Cell type enriched (Skin - Keratinocyte (other), Testis - Spermatogonia) |
| Immune cell specificity | Low immune cell specificity |
| Immune cell expression cluster | Non-specific - Cell proliferation (mainly) |
| Prognostic summary | SLC35F2 is a prognostic marker in Lung adenocarcinoma, Lung squamous cell carcinoma, Pancreatic adenocarcinoma |
| Cancer specificity | Low cancer specificity |
| Cell line expression cluster | Non-specific - Unknown function (mainly) |
| Cell line specificity | Low cancer specificity |

Top 15 nearest neighbours based on tissue RNA expression

| Neighbour | Description | Correlation | Cluster |
|---|---|---|---|
| GK5 | Glycerol kinase 5 | 0.8544 | 45 |
| BTC | Betacellulin | 0.8544 | 45 |
| GOLPH3L | Golgi phosphoprotein 3 like | 0.8088 | 24 |
| GGT6 | Gamma-glutamyltransferase 6 | 0.8070 | 48 |
| SP8 | Sp8 transcription factor | 0.8035 | 64 |
| HOXB13 | Homeobox B13 | 0.7965 | 45 |
| ESRP1 | Epithelial splicing regulatory protein 1 | 0.7947 | 48 |
| CWH43 | Cell wall biogenesis 43 C-terminal homolog | 0.7947 | 27 |
| FXYD3 | FXYD domain containing ion transport regulator 3 | 0.7772 | 45 |
| STX19 | Syntaxin 19 | 0.7772 | 53 |
| INAVA | Innate immunity activator | 0.7649 | 45 |
| OR2A7 | Olfactory receptor family 2 subfamily A member 7 | 0.7632 | 45 |
| ZNF587 | Zinc finger protein 587 | 0.7632 | 48 |
| SHROOM3 | Shroom family member 3 | 0.7632 | 75 |
| CDCP1 | CUB domain containing protein 1 | 0.7614 | 75 |

Cluster 48: Stomach - Proteolysis
Cluster 45: Intestine - Digestion

Summary of SLC35F2 co-expressing genes (source: https://seek.princeton.edu/seek/)

Top 10 SLC35F2 co-expressing genes in top weighted datasets

| Rank | Z-score of Pearson correlation | Gene |
|---|---|---|
| 1 | 1.742 | SPIN4 |
| 2 | 1.715 | SFN |
| 3 | 1.686 | SNHG1 |
| 4 | 1.682 | EPCAM |
| 5 | 1.670 | SPINT1 |
| 6 | 1.646 | EHF |
| 7 | 1.625 | F2RL1 |
| 8 | 1.623 | CHMP4C |
| 9 | 1.615 | DSG2 |
| 10 | 1.593 | CLDN4 |

Gene set enrichment analysis (GO (BP) term) of top 100 SLC35F2 co-expressing genes

| Term | p-value ? | q-value ? |
|---|---|---|
| protein complex disassembly | 7.09E-8 | 1.07E-6 |
| cellular protein complex disassembly | 7.09E-8 | 1.44E-6 |
| translational termination | 7.09E-8 | 1.44E-6 |
| viral transcription | 2.11E-7 | 4.45E-6 |
| cellular component disassembly | 1.47E-5 | 1.43E-4 |
| establishment of protein localization to organelle | 7.85E-5 | 7.56E-4 |
| interspecies interaction between organisms | 2.88E-4 | 1.99E-3 |
| translation | 3.02E-4 | 3.03E-3 |
| viral process | 5.26E-4 | 4.15E-3 |
| single organism membrane organization | 5.7E-4 | 4.29E-3 |
| protein targeting | 5.7E-4 | 4.3E-3 |
| keratinocyte proliferation | 8.39E-4 | 6.35E-3 |
| regulation of ventricular cardiac muscle cell action potential | 8.39E-4 | 9.71E-3 |
| release of cytochrome c from mitochondria | 4.67E-3 | 8.3E-2 |
| ribosomal small subunit biogenesis | 1.7E-2 | 1.38E-1 |
| actin filament based process | 1.7E-2 | 1.61E-1 |

**Figure S2. Co-expression analyses link the human SLC35F2 but not the human SLC35F1 transporter to ribosome biogenesis and RNA processing.** Co-expression data from the Protein atlas (https://www.proteinatlas.org/) and SEEK (https://seek.princeton.edu/seek/) databases for SLC35F1 **(A)** and SLC35F2 **(B)**.

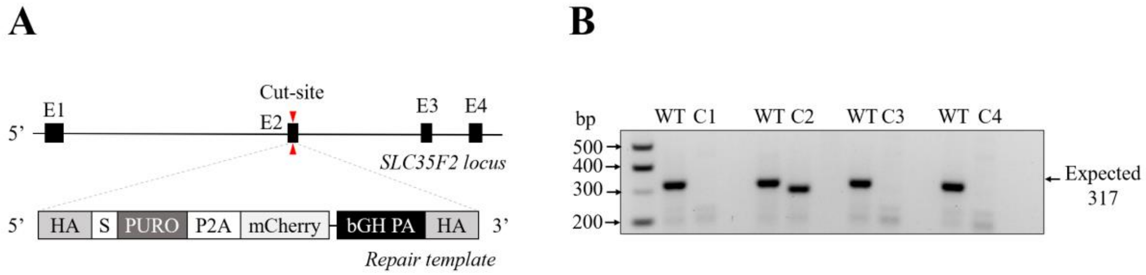


**Figure S3. Targeting of the SLC35F2 locus in HeLa cells. (A)** The SLC35F2 gene, located on the reverse strand of Chromosome 11 in humans, was disrupted through the insertion of a PURO-P2A-mCherry selection cassette into the second coding exon (E2) of the gene. **(B)** Screening of four SLC35F2 candidate targeted clones (C1-C4) via PCR. Amplicons from non-targeted HeLa cells represent positive controls and show the wild-type SLC35F2 allele (WT), which is 317 bp (+) in length.

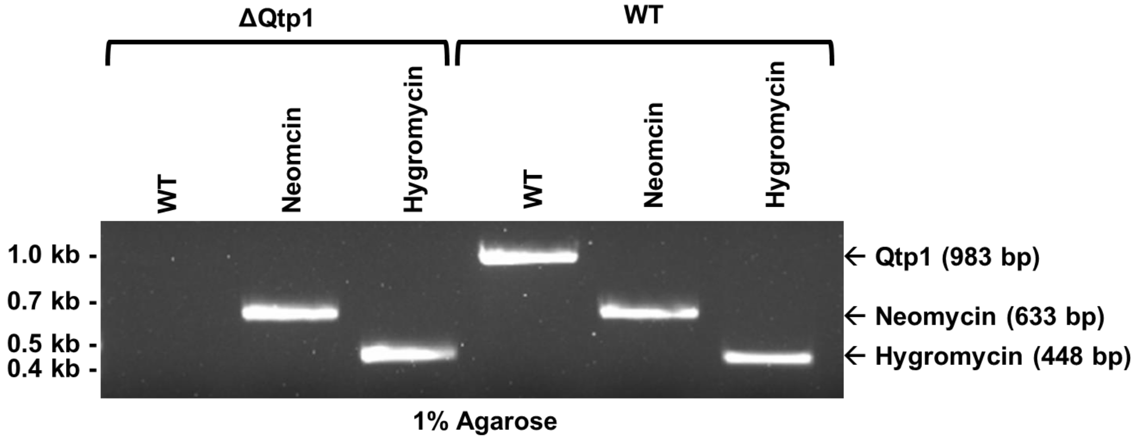


**Figure S4**. PCR confirmation of Qtp1 double deletion in *T. brucei*. Amplification of the ORFof Qtp1 from isolated gDNA does not generate products in the deletion strain.

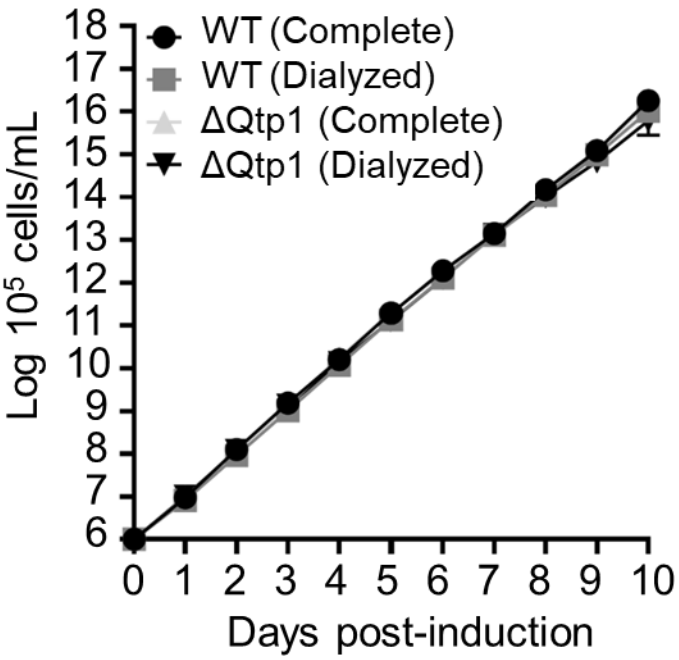


**Figure S5.** Supplementation of queuosine and queuine to 25 nM final concentration after depletion. Triplicate growth curve of *T. brucei* cells comparing growth between WT and ΔQtp1 strains cultured in media supplemented with either complete or dialyzed FBS.

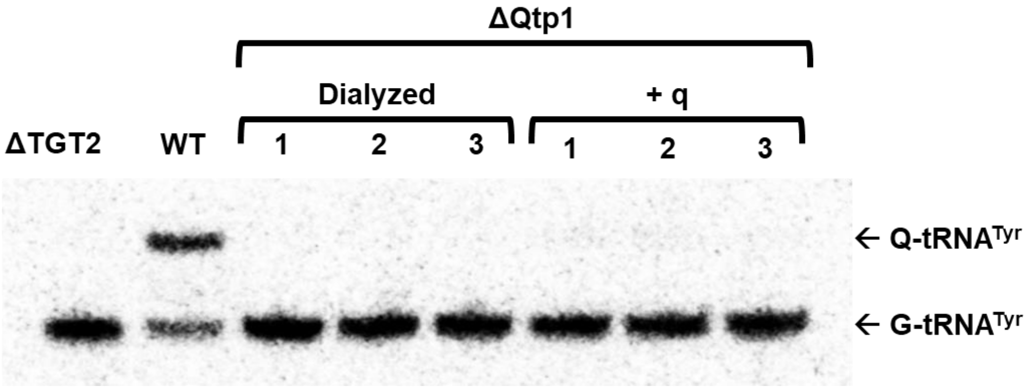


**Figure S6**. Supplementation of queuine to 250 nM final concentration after depletion in *T. brucei*. APB-Northern blot hybridization performed on extracted total RNA samples revealing both Q-$tRNA^{Tyr}$ and G-$tRNA^{Tyr}$.

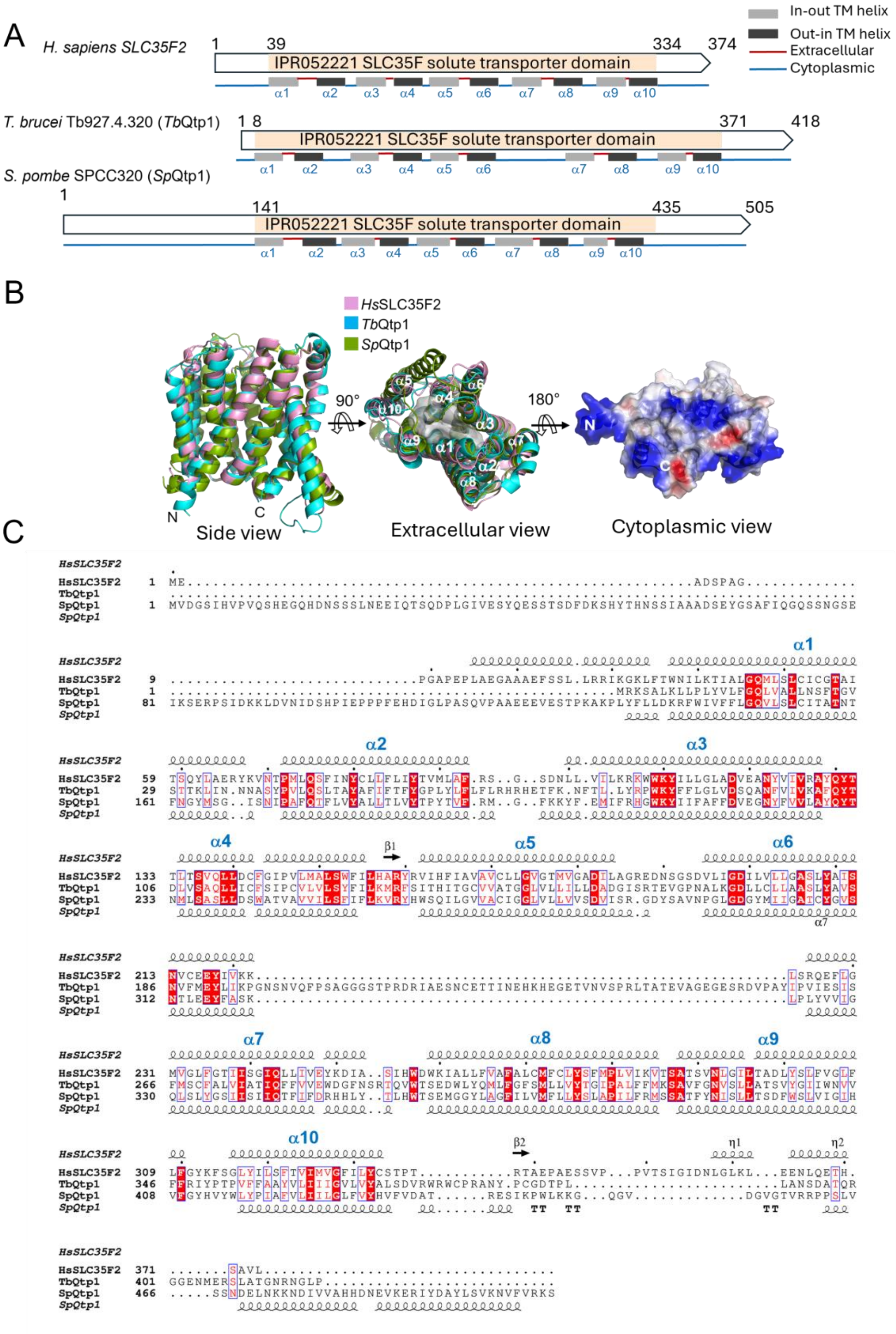
A
H. sapiens SLC35F2
IPR052221 SLC35F solute transporter domain
1
39
334
374
In-out TM helix
Out-in TM helix
Extracellular
Cytoplasmic
T. brucei Tb927.4.320 (TbQtp1)
1 8
371
418
S. pombe SPCC320 (SpQtp1)
141
435
505
α1
α2
α3
α4
α5
α6
α7
α8
α9
α10
B
HsSLC35F2
TbQtp1
SpQtp1
90°
180°
N
C
Side view
Extracellular view
Cytoplasmic view
C
HsSLC35F2
TbQtp1
SpQtp1

**Figure S7. Structural view of SLC35F2/Qtp1 proteins. (A)** Schematic of the primary structures of human SLC35F2, *T. brucei* Qtp1 and *S. pombe* Qtp1 proteins. The core transporter domain in each protein and its ten predicted transmembrane helices (10-TM) are indicated. The color code indicates helix orientation in the membrane and intracellular versus extracellular regions, as predicted by Deep TMHMM. **(B)** Superposed AlphaFold-predicted models of the three proteins inside (left) and extracellular (middle) views with the transmembrane helices labeled and central binding site cavity shown as a grey surface. For clarity, the N- and C-terminal regions outside the transporter 10-TM domain, and a 60-residue unstructured region between helices $\alpha$6 and $\alpha$7 in *T. brucei* Qtp1 are not shown. Right: Electrostatic surface potential showing positively charged surface of the putative cytoplasmic side which harbors the N- and C-termini of the core transporter domain. **(C)** Structure-based multi-sequence alignment generated from the aligned models. Predicted secondary structure elements from human SLC35F2 and *S. pombe* Qtp1 are shown above and below the sequences. *Hs*: *H. sapiens*, *Tb*: *T. brucei*, *Sp*: *S. pombe*.

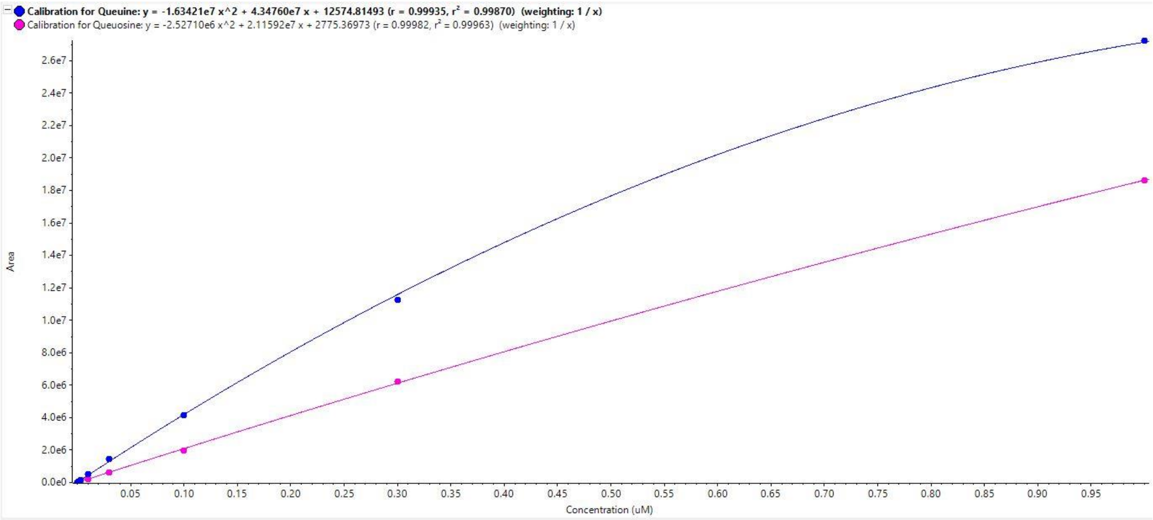


**Figure S8.** Calibration curves used to calculate the extracellular (Media) and intracellular (Cytosol) levels of queuine (blue line) and queuosine (pink line) in Fig. 4. Plotted are the MS peak areas vs. concentration (μM) using 8 calibrators for queuine and queuosine standards. Line equations and regression fitting values are found at the top of the x-axis.

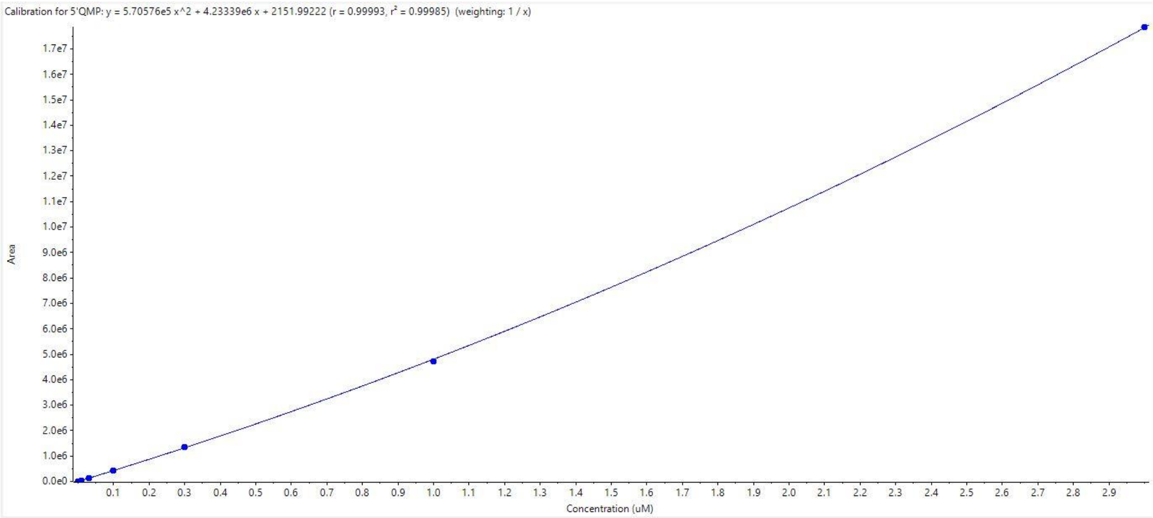


**Figure S9.** Calibration curves used to calculate the extracellular (Media) and intracellular (Cytosol) levels of Q-5′MP (blue line) in Fig. 4. Plotted are the MS peak areas vs. concentration (µM) using 8 calibrators for Q-5′MP standards. Line equation and regression fitting values are found at the top of the x-axis.
